# MCRS1 modulates the heterogeneity of microtubule minus-end morphologies in mitotic spindles

**DOI:** 10.1101/2022.06.03.494715

**Authors:** Alejandra Laguillo-Diego, Robert Kiewisz, Carlos Martí-Gómez, Daniel Baum, Thomas Müller-Reichert, Isabelle Vernos

## Abstract

Faithful chromosome segregation requires the assembly of a bipolar spindle, consisting of two antiparallel microtubule (MT) arrays having most of their minus ends focused at the spindle poles and their plus ends overlapping in the spindle midzone. Spindle assembly, chromosome alignment and segregation require highly dynamic MTs. The plus ends of MTs have been extensively investigated; instead, their minus end structure remains poorly characterized. Here, we used large-scale electron tomography to study the morphology of the MT minus ends in 3D-reconstructed metaphase spindles in HeLa cells. In contrast to the homogeneous open morphology of the MT plus ends at the kinetochores, we found that MT minus ends are heterogeneous showing either open or closed morphologies. Silencing the minus-end specific stabilizer, MCRS1 increased the proportion of open MT minus ends. Altogether, these data suggest a correlation between the morphology and the dynamic state of the MT ends. Taking this heterogeneity of the MT minus end morphologies into account, our work indicates an unsynchronized behavior of MTs at the spindle poles, thus laying the ground for further studies on the complexity of MT dynamics regulation.

## Introduction

The mitotic spindle is a highly complex molecular machinery that assembles during cell division to segregate chromosomes to the daughter cells. It is structurally and functionally defined by its main component, the microtubules (MTs). During mitosis, MTs are nucleated through different pathways, involving the centrosomes, the chromosomes and pre-existing MTs (Lüders, 2016; Meunier and Vernos, 2012; Teixido-Travesa et al., 2012). Altogether, the bipolar spindle consists of MTs with their plus ends at the spindle equator and the minus ends associated with the two spindle poles (i.e., the centrosomes) or with other MT lattices (Kiewisz et al., 2022). Within the spindle of mammalian cells, kinetochore MTs (KMTs) organize into bundles, so-called kinetochore-fibers (k-fibers) that connect chromosomes to the two spindle poles. In addition, astral (AMTs) and interpolar MTs (IMTs; both termed non-KMTs here) constitute the main body of the spindle, providing additional forces for spindle positioning and chromosome movements by interaction with molecular motors (Meunier and Vernos, 2012; Meunier and Vernos, 2016).

The regulation of MT dynamics is fundamental for the overall organization of the spindle and the segregation of the chromosomes. Dynamics can be observed at both MT ends that can stochastically switch between phases of growth and shrinkage (Kirschner and Mitchison, 1986; Mitchison and Kirschner, 1984). MT plus-end dynamics promotes the attachment of MTs to the kinetochores and is important for the assembly of the k-fibers (Mimori-Kiyosue and Tsukita, 2003). The dynamic nature of these attachments is key for error correction, the activity of the spindle assembly checkpoint (SAC) and chromosome segregation (Foley and Kapoor, 2013; Lampson and Grishchuk, 2017; Maiato et al., 2004; Monda and Cheeseman, 2018). Although MT minus-end dynamics seem to play a role in spindle assembly and chromosome movements, little information is currently available on the structure and dynamics of the MT minus ends in the mammalian spindle, as most of them are focused at the spindle poles, an extremely crowded region difficult to observe and manipulate (Akhmanova and Steinmetz, 2019).

In mammalian cells, most MTs are nucleated by the γ-TuRC, a dedicated multiprotein complex. Although MTs in mitosis are nucleated through various pathways, spindle pole-located γ-TuRC plays a crucial role in MT nucleation. In general, MTs nucleated from γ-TuRC are “capped” at their minus ends, and they are therefore stabilized (Consolati et al., 2020; Wieczorek et al., 2020; Zimmermann et al., 2020). However, various studies indicate that MT minus ends depolymerize at the spindle poles (Jiang et al., 2017). Indeed, a movement of tubulin subunits from the spindle equator to the spindle poles was observed by live-cell imaging upon photoactivation of a tubulin stripe in close proximity to the aligned chromosomes in the metaphase spindle (Mitchison, 1989). This phenomenon, named spindle flux, was proposed to play a role in the control of spindle length and in the chromosome segregation (Ganem and Compton, 2006; Ganem et al., 2005; Steblyanko et al., 2020). The current view is that the spindle flux is the result of a combination of MT transport towards the spindle poles and net incorporation of tubulin at the MT plus ends. This incorporation is compensated by the removal of tubulin dimers at MT minus ends at the spindle poles (Kirschner and Mitchison, 1986; Mitchison, 1989; Waters et al., 1996) caused by kinesin-13 depolymerases (Ems-McClung and Walczak, 2010). Consistently, silencing the spindle pole-localized kinesin-13 depolymerase, kif2a leads to the formation of monopolar spindles, indicating that the regulation of MT minus end dynamics is essential for bipolar spindle assembly (Ganem and Compton, 2004). Additional support for this idea was also provided by the identification of Microspherule Protein 1 (MCRS1), a RanGTP-regulated protein that localizes to the KMT minus ends and regulates their depolymerization rate at the spindle poles. Spindles assembled in MCRS1-silenced cells have a faster poleward flux, show hyper-stretched kinetochores and form unstable spindles (Meunier and Vernos, 2011).

Several studies indicate that the dynamic state of MTs is associated with specific end morphologies. Cryo-electron microscopy studies of *in vitro*-assembled MTs showed that fast growing MTs have flared ends with curved sheet-like protofilaments at their tips, while slowly growing MTs have blunt ends (Chretien et al., 1995; Mandelkow et al., 1991; Muller-Reichert et al., 1998; Rice, 2018; Simon and Salmon, 1990). In contrast, depolymerizing MT ends display outward-curled (also called ramshorn-like) protofilaments (Chretien et al., 1995; Mandelkow et al., 1991; Muller-Reichert et al., 1998; Rice, 2018; Simon and Salmon, 1990). Importantly, closed MT ends have also been observed. MTs nucleated by the yTuRC complex *in vitro* show an electron-dense material, thus ‘closing’ their minus-ends (Zheng et al., 1995).

Although MT dynamics in cells is more complex than *in vitro* due to the presence of a large variety of MT-associated proteins (MAPs), including some that specifically bind to the MT ends, the morphologies of both growing and depolymerizing ends were found to be very similar to those observed *in vitro* (VandenBeldt et al., 2006). Moreover, recent electron microscopy studies of plastic-embedded spindles in *C. elegans* also revealed flared and curled MT end morphologies (O’Toole et al., 2003), suggesting that the spindle MTs may be in different phases of polymerization and depolymerization, respectively. MTs having a blunt end may be pausing, polymerizing or depolymerizing, and therefore it is difficult to categorize them in any specific dynamic state. However, more recent tomographic reconstructions *in vitro* and in cells display similar bent MT tips in growing and shortening MTs, indicating that dynamic states cannot be distinguished that easily (Gudimchuk and McIntosh, 2021; Gudimchuk et al., 2020; McIntosh et al., 2018).

Interestingly, partial reconstructions of metaphase spindles in U2OS cells displayed a mixture of closed and open morphologies at the pole-facing MT ends (i.e., at the putative minus ends; (Kamasaki et al., 2013). Consistently, studies performed in early *C. elegans* embryos showed that the minus ends of KMTs have heterogeneous morphologies described as either closed or open (O’Toole et al., 2003). The presence of a “cap-like” structure at the MT minus ends associated with either spindle pole bodies in *S. pombe* or mitotic centrosomes in *C. elegans* embryos was interpreted as the yTuRC (Hoog et al., 2011; O’Toole et al., 2003; Teixido-Travesa et al., 2012). However, a systematic characterization of MT minus ends in mitotic spindle poles of mammalian cells is currently not available.

Here, we set out to gain novel insights into the MT minus-end morphologies in the metaphase spindle of human cells using electron tomography. We found that MT minus ends can have either closed or open morphologies in a proportion that is modified upon silencing by the MT minus end regulator MCRS1. The observed heterogeneity of KMT minus-end morphologies suggests a complex mechanism of regulation of their dynamics.

## Results

### Different MT minus-end morphologies co-exist at the spindle poles

We used 3D reconstructions of whole mitotic spindles assembled in HeLa cells (Kiewisz et al., 2022) to analyze the morphology of the MT ends (**Fig. 1A**, **Fig S1A, C, E, G** and **Video 1**; **Table 1–2**). In these control cells (called ‘control’ hereafter), we defined two main classes of ends: closed ends that have an electron-dense “cap” (**Fig. 1B**, left panels; **Fig. 1, Fig. S5A**) and open ends that typically have a flared morphology (**Fig. 1B**, right panels; **Fig. S5B**). Other types of MT ends were classified as ‘undefined’.

**Figure 1.**
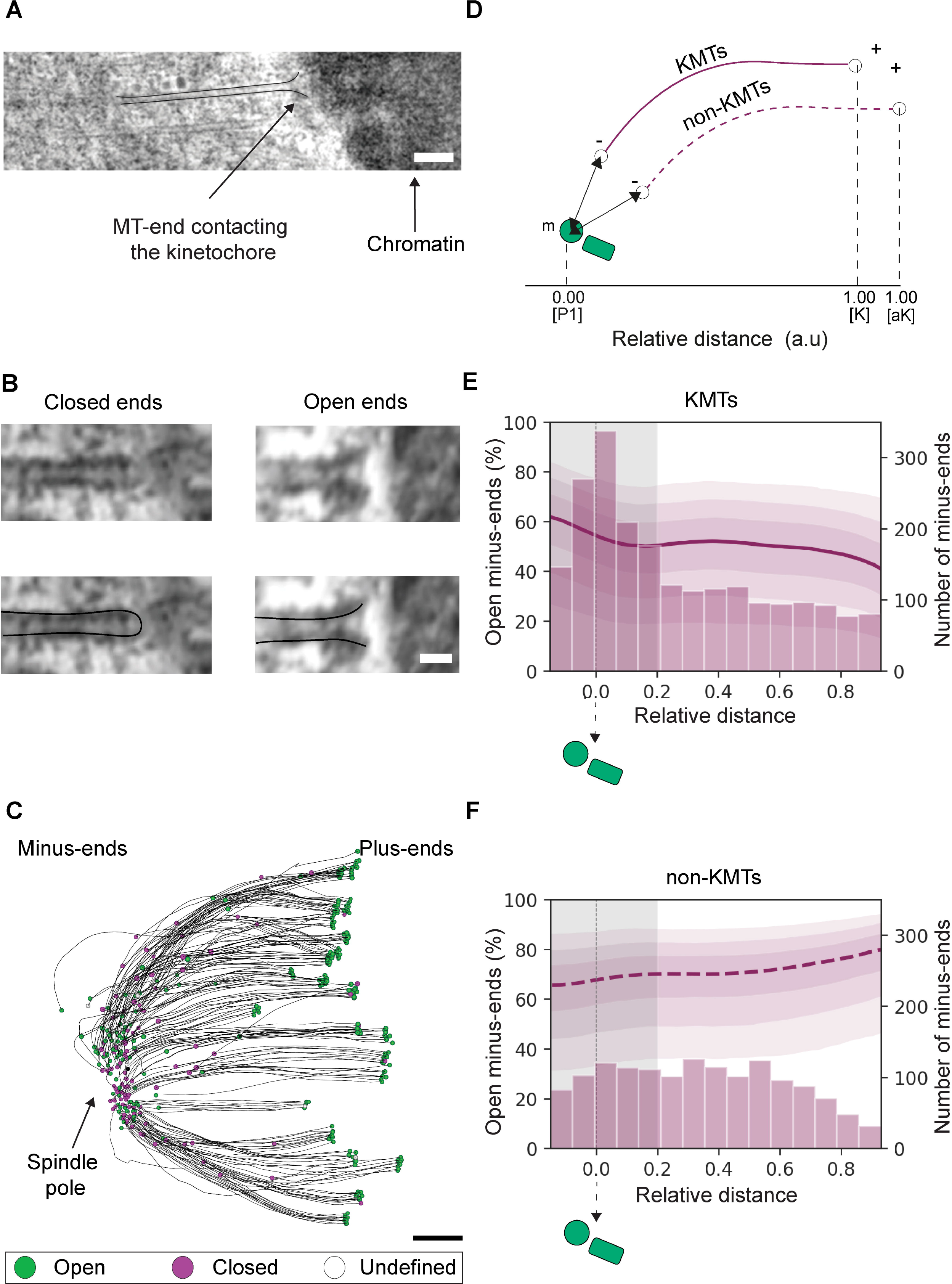
Quantitative analysis of MT minus-end morphologies. (**A**) Tomographic slice showing associated with a chromosome (arrow) as visualized in a HeLa cell at metaphase. In this study, such an MT is defined as a KMT (top view, tomographic slice with overlay). Scale bar, 100 nm. (**B**) Visualization of a closed and an open MT end morphology (top row, tomographic slice; bottom row, tomographic slices with overlays for better illustration). MT ends showing an electron-dense cap are classified as closed (left panels). Curved or sheet-like ends are classified as open (right panels). Scale bar, 25 nm. (**C**) Orthogonal projection (top view) of a 3D model showing a quarter volume of a control spindle (Ctrl (#3)). KMTs are depicted as black lines and their end morphologies are indicated as open (green circles), closed (purple circles) or undefined (open circles). The position of the spindle pole is indicated by an arrow. Scale bar, 500 nm. (**D**) Schematic drawing illustrating the positional analysis of both KMT and non-KMT minus ends. The position of each MT minus end along the spindle pole axis is given as a relative distance. The relative distance of each KMT minus end along the half-spindle axis is defined by the position between the pole [P1, 0] and the kinetochore [K, 1]. The relative position of non-KMT minus ends is defined by the position of the pole and the average position of all kinetochores in the spindle [aK, 1]. The absolute distance (arrows) is determined by measuring the 3D distance of each MT minus end to the nearest mother centriole (m). A KMT is indicated by a solid purple line, a non-KMT by a dashed purple line. (**E**) Histogram showing the absolute number of KMT minus ends (right axis) at different relative distances. The graph also shows the estimated percentage of open minus ends at each given relative position (left axis). The different levels of shadow around the line (purple) correspond to the credible intervals (CI), representing 95, 80 and 50% probability to observe the true value within these intervals. The shaded area (grey) represents the MT-spindle pole association area. This area was defined as the region of the spindle pole with the highest concentration of KMT minus ends as observed in the histogram. The total number of classified KMT minus ends for control spindles was n = 2071. (**F**) Histogram showing the absolute number of non-KMT minus-ends in the MT spindle pole association area. The total number of classified non-KMT minus-ends is n = 1516.

**Table 1.** Tomographic data sets as used throughout this study.

| Metadata/<br>data set | Original<br>data set | Spindle<br>reconstruction | No. of serial<br>sections<sup>1</sup>/<br>Z-factor | Estimated<br>tomographic volume<br>($\mu\text{m}^3$) | Data<br>size<br>(Gb) |
| --- | --- | --- | --- | --- | --- |
| Ctrl #1 | T_0475 | Full<br>(divided into<br>quarters) | 22 / 1.30 | 144 | 44.3 |
| Ctrl #2 | T_0479 | Full<br>(divided into<br>quarters) | 29 / 1.40 | 263 | 74.2 |
| Ctrl #3 | T_0550_wt_1 | Quarter | 17 / 1.30 | 114 | 30 |
| siScr #1 | T_0506 | Quarter | 12 / 1.25 | 58 | 22.4 |
| siScr #2 | T_0550_scr_3 | Quarter | 14 / 1.30 | 78 | 28.3 |
| siMCRS1<br>#1 | T_0506_mcrs1_2 | Quarter | 18 / 1.32 | 94 | 26.8 |
| siMCRS1<br>#2 | T_0506_mcrs1_4 | Quarter | 18 / 1.35 | 87 | 24.9 |
| siMCRS1<br>#3 | T_0506_mcrs1_5 | Quarter | 17 / 1.30 | 79 | 22.8 |
| siMCRS1<br>#4 | T_0550_mcrs1_2 | Quarter | 16 / 1.35 | 112 | 32.1 |
| siMCRS1<br>#5 | T_0550_mcrs1_3 | Quarter | 21 / 1.36 | 110 | 31.6 |

<sup>1</sup> Average section thickness is 300 nm.

**Table 2.** Quantitative analysis of the tomographic reconstructions as used throughout this study.

| Data set | No. of MTs | No. of KMTs | No. of KMTs in the reconstructed volume | No. of k-fibers | Pole-to-pole distance (μm) | Distance between outer kinetochores (μm) |
| --- | --- | --- | --- | --- | --- | --- |
| Ctrl #1<br>(Kiewisz et al., 2022) | 4884 | 797 | 748 | 92 | 7.16 | 1.08 ±0.20<br>(n=43) |
| Ctrl #2<br>(Kiewisz et al., 2022) | 8047 | 1102 | 1072 | 110 | 10.39 | 1.24 ±0.21<br>(n=50) |
| Ctrl #3 | - | 407 | 205 | 22 | - | 1.15 ±0.22<br>(n=22) |
| siScr #1 | 2913 | 397 | 203 | 20 | - | 1.02 ±0.20<br>(n=17) |
| siScr #2 | 6221 | 344 | 205 | 24 | - | 1.06 ±0.21<br>(n=28) |
| siMCRS1 #1 | 2884 | 224 | 50 | 8 | - | 1.10 ±0.24<br>(n=20) |
| siMCRS1 #2 | 2451 | 169 | 57 | 10 | - | 1.12±0.23<br>(n=10) |
| siMCRS1 #3 | 3635 | 301 | 159 | 18 | - | 1.06 ±0.11<br>(n=6) |
| siMCRS1 #4 | 5624 | 348 | 180 | 29 | - | 1.02 ±0.18<br>(n=28) |
| siMCRS1 #5 | 3811 | 484 | 273 | 39 | - | 1.13±0.20<br>(n=35) |

**Table 3.** Quantitative analysis of MT minus-end morphologies

| End morphology | KMT minus ends |  |  |  | KMT plus ends |  |  |  |
| --- | --- | --- | --- | --- | --- | --- | --- | --- |
|  | Total no. | Matching open ends | Matching closed ends | Undefined ends | Total no. | Matching open ends | Matching closed ends | Undefined ends |
| **Control**<br>(Ctrl #1, #2, #3) | 2239 | 682 | 385 | 532 | 1841 | 1509 | 6 | 218 |
| **siScramble**<br>(siScr #1, #2) | 407 | 109 | 66 | 137 | 432 | 350 | 2 | 47 |
| **siMCRS1**<br>(siMCRS1 #1, #2, #3, #4, #5) | 726 | 297 | 90 | 185 | 847 | 692 | 2 | 108 |
| End morphology | Non-KMT minus ends |  |  |  |  |  |  |  |
|  | Total no. of ends | Matching open ends | Matching closed ends | Undefined ends |  |  |  |  |
| **Control**<br>(Ctrl #1, #2, #3) | 1511 | 521 | 209 | 427 |  |  |  |  |
| **siScramble**<br>(siScr #1, #2) | 1155 | 278 | 59 | 591 |  |  |  |  |
| **siMCRS1**<br>(siMCRS1 #1, #2, #3, #4, #5) | 6654 | 1876 | 509 | 3539 |  |  |  |  |

We first focused on the MTs forming the k-fibers (i.e., the KMTs), strictly defined as those directly associated through their plus ends to the outer kinetochores at the chromosomes (Kiewisz et al., 2022). In agreement with the dynamic nature of the KMT plus ends (Cheeseman and Desai, 2008), the close examination of the KMT end morphology at the kinetochore showed that they were open (green circles) and flared (**Fig. 1B**– right panel and **Fig. 1C**) (O’Toole et al., 2003). We then focused on the opposite end of the KMTs away from the kinetochore defining this one as the minus end. KMT minus ends had either closed (**Fig. 1C**, circles in purple) or open morphologies (**Fig. 1C**, circles in green). In the context of previous work on MT minus-end dynamics and morphology (Hoog et al., 2011), this suggested to us that the heterogeneous morphologies of the KMT minus ends reflect the co-existence of different dynamic states.

Previous work had indicated that about half of the KMT minus ends are located at the spindle poles (defined as the MT-centrosome interaction area), while the other half is distributed along the half-spindle (Kiewisz et al., 2022). Extending this previous analysis, we were also interested to know whether there was a correlation between the KMT minus-end morphologies and the positioning in the spindle. For this, we plotted the distribution of KMT minus ends relative position on the kinetochore-to-spindle axis (**Fig. 1D**). The kinetochore-to-spindle axis was defined as the distance between the closest spindle pole (position = 0) and the individual kinetochores (position = 1). Additionally, we also defined the MT-centrosome interaction area as twice the half-width. We then determined the proportion of open minus ends at relative distances along the kinetochore-to-spindle axis (**Fig. 1E**). The inferred proportion of the open minus ends in the proximity of the centriole pair (defining the spindle pole) was 52.97% (95% credible interval (CI) = [21.48, 77.43]%) (**Fig. 1E,** gray vertical area D < 0.2, region where the majority of KMT minus ends are located). Therefore, about half of the KMT minus ends at the spindle poles were open, suggesting that they are in different dynamic states.

We then asked whether KMTs belonging to the same k-fiber might have similar minus end morphologies indicative of a synchronized dynamic state. To address this question, we performed a model comparison using a likelihood ratio test between models that included or ignored the k-fiber as a random effect. We found no evidence of clustering of open minus ends at specific k-fibers (p-value = 1). This suggested that the minus-end morphology of any given KMT within a k-fiber is independent from the minus end morphologies of the other KMTs in the same k-fiber.

To determine whether the mixture of minus end morphologies suggestive of mixed dynamic states is specific for the KMTs or a more general feature of the spindle MTs, we extended our studies to the non-KMTs. For these MTs, we defined the average position of all kinetochores (**Fig. 1D,** aK) as position 1 on the half-spindle axis. In addition, the polarity of the non-KMTs was determined by the positioning of the ends along the half-spindle axis. The end of each non-KMT closer to the spindle pole was defined as the minus end, with the other end assigned as the plus end. The end morphologies of non-KMTs were also classified as either closed or open. We then estimated the proportion of open minus ends at each relative distance. We found that 64.04% (CI = [33.00, 86.22]%) of the non-KMT minus ends were open in the region near the centriole pair (**Fig. 1F**, gray area). Again, these data showed that MT minus ends have heterogeneous morphologies, overall suggesting the coexistence of different dynamic states.

### MCRS1 silencing induces changes in k-fiber ultrastructure and spindle shape

MCRS1 was shown to associate with the k-fiber minus ends in metaphase and regulate their dynamics (Meunier and Vernos, 2011). Therefore, we decided to examine the changes in the MT end morphologies in spindles assembled in MCRS1-silenced cells by electron tomography. Since siRNA-based gene silencing may not be homogeneous in a cell population, we first carefully looked for morphological spindle features that could be altered in MCRS1-silenced cells and used as a signature for selecting the spindles to be processed for electron tomography. Western blot analysis of MCRS1-silenced cells showed a reduction in the target protein level of close to 90% (**Fig. 2A-B and Fig. S3**). High-contrast light microscopy images revealed specific changes in the half-spindle shape and the outer spindle MT angle (**Fig. 2C** and **Fig. S4**). siScramble cells (i.e.: transfected with random siRNA) showed an average half-spindle angle of 80.81° ±10.60 (**Fig. 2E**, n=105). In contrast, MCRS1-silenced cells had an average half-spindle angle of 71.5° ±8.05 (n=110; p-value = 4.729 × 10^−8^), significantly narrower than in siScramble cells. In addition, siScramble cells showed an average angle for the outer MTs of 152.34° ±9.62 (**Fig. 2G**, n=201,), whereas siMCRS1-depleted cells displayed an average angle for the outer MTs of 167.51° ±10.72 (n=161, p-value = 7.486 × 10^−26^). Thus, the outer MTs in MCRS1-depleted cells were significantly straighter than in siScramble cells. Similar differences were also clearly observed in 3D models after processing both siScramble and MCRS1-silenced cells for electron tomography. In stacked serial plastic sections (**Fig. 2D**), siScramble cells had a half-spindle angle of 90.62° ±6.41 (**Fig. 2F**, **Video 2**; n=4), whereas MCRS1-silenced spindles had a significantly lower half-spindle angle of 80.13° ±5.77 (**Video 3**; n=8, p-value=0.017). Moreover, siScramble spindles showed an average outer MT angle of 145.77° ±5.55 (**Fig. 2H, Video 3**; n=8,), whereas siMCRS1 spindles displayed a significantly higher average outer MT angle of 158.00° ±8.74 (n=16, p-value = 0.002). From these data, we concluded that the depletion of MCRS1 caused a reduction in the half-spindle angle and a simultaneous increase in the MT angle.

**Figure 2.**
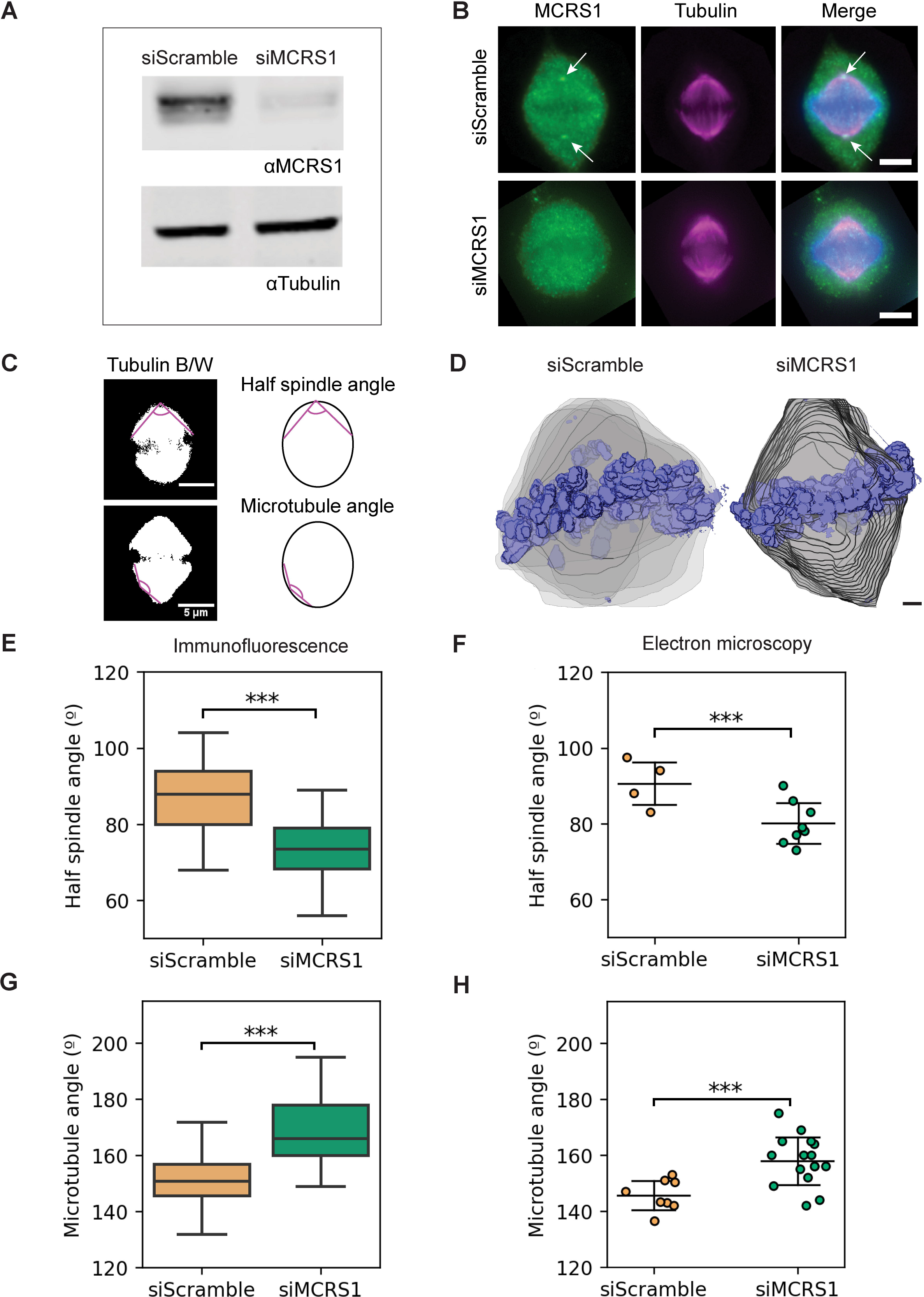
Characterization of metaphase spindles in MCRS1-silenced cells. (**A**) Western Blot analysis showing the levels of MCRS1 in control and MCRS1-silenced cells. Tubulin levels were used as a loading control. (**B**) Immunofluorescence staining of siScramble (top panel) and MCRS1-silenced cells (lower panel). MCRS1 is shown in green, DNA in blue and tubulin in magenta. MCRS1 staining can be observed as tiny spots at spindle poles in the control cell (arrows). Upon silencing, MCRS1 cannot be detected anymore. Scale bars, 5 μm. (**C**) Black/white images of control (top, left) and a silenced spindle (bottom, left). The magenta lines show the angles that were measured for the half spindle (top, right) and for the MTs in the outward position of the spindle (bottom, right). Scale bars, 5 μm. (**D**) Three-dimensional models of spindle shape in siScramble (#2) and siMCRS1 (#5) cells were obtained from low-resolution screening by transmission electron microscopy. The outline of the spindle volume in each serial section is shown in gray and chromosomes in blue. Scale bar, 1 μm. (**E**) Boxplot showing the quantification of the half-spindle angle obtained from immunofluorescence images. The boxes show the upper and lower quartiles, the whiskers show the minimum and maximal values excluding outliers; the line inside the box indicates the median. Asterisks show a significant difference according to a linear regression model considering two different experiments with a p-value=4.729 × 10^−8^ (siScramble: n=105; siMCRS1: n=110). (**F**) Scatterplot showing the quantification of the half-spindle angle from electron microscopic images. The line shows the mean, and the error bars represent ±SD. Asterisks show significant differences according to a two-tailed student t-test with a p-value=0.017. (siScramble: n=4; siMCRS1: n=8). (**G**) Boxplot showing the quantification of the half-spindle angle obtained from immunofluorescence images. Asterisks show a significant difference according to a linear regression model considering two different experiments with a p-value=7.486 × 10^−26^ (siScramble: n=201; siMCRS1: n=161). (**H**) Scatterplot showing the quantification of the MT angle obtained from electron microscopic images (two-tailed student t-test with a p-value = 0.002 (siScramble: n=8; siMCRS1: n=16).

These parameters were then used for the selection of spindles from siScramble cells and MCRS1-silenced cells for 3D reconstruction. To obtain representative data in particular for spindles assembled in MCRS1-silenced cells, we aimed at obtaining data from several different spindles (**Video 4-5**). We then evaluated whether reconstructions of a quarter of the spindle volume would be representative of a full spindle. Using the tomograms from control samples, we divided a full spindle into 4 symmetrical quarters and measured the number of KMTs per k-fiber and the outer-kinetochore distance (**Fig. S6–7**). We found no differences between these values and those obtained from the full spindle tomogram analysis. This suggested that the quarter spindles are representative of the corresponding full spindle. Therefore, we decided to reconstruct quarters from two spindles from siScramble cells and five from MCRS1-silenced cells, which allowed us to increase the number of analyzed individual data sets. To determine whether all the selected spindles were at a similar stage in mitosis, we compared the average distances between the outer-kinetochores of sister k-fibers for all our data sets. The average inter-kinetochore distance in the selected spindles was 1.06 ±0.21 μm (n=28) in siScramble cells and 1.06 ±0.21 μm (n=55) in MCRS1-silenced cells. Control spindles showed an average value of 1.07 ±0.21 μm (n=146; siScramble *versus* Control, p-value=0.108; siScramble *versus* siMCRS1, p-value=0.240). Additionally, the values measured for Control and siScramble cells were in agreement with previous studies (Kiewisz et al., 2022). The high similarity in the inter-kinetochore distance in all these conditions indicated that the selected spindles were captured at a similar mitotic stage.

We then analyzed the 3D reconstructions of all spindles. MCRS1-silenced spindles showed distinct ultrastructural features. The k-fibers appeared straighter in siMCRS1 compared to siScramble spindles (**Fig. 3A, Fig. S2, Fig. S8; Video 6-7** and **also see Video 4-6** from (Kiewisz et al., 2022)). Interestingly, the KMTs did not reach the centriole pair area at the spindle poles in siMCRS1 spindles, whereas the centriole pair was embedded within the mass of KMT minus ends in the siScramble spindles. Consistently, the quantification of MT minus end distribution revealed that the majority reached the centrioles in control (**Fig. 3B**, mean peak at position = 0.03) and siScramble spindles (**Fig. 3B**, mean peak at position = 0.04, p-value=0.201). In contrast, the MT minus ends in siMCRS1 spindles peaked at a mean relative position of 0.07, further away from the centrioles (**Fig. 3B**, p-value = 0.018; *siScramble vs siMCRS1 at D < 0.2*). The displacement of MT minus ends away from the spindle pole could result from a higher MT minus-end depolymerization which would be consistent with the proposed role of MCRS1 in controlling the rate of MT minus end depolymerization. We then determined the number of KMTs per k-fiber in the three different conditions (**Fig. 3C**) and evaluated the putative differences using a Generalized Linear Model (GLM) with Poisson likelihood and log link function, using siMCRS1 and siScramble data as covariates. We found that the number of KMTs per k-fiber was reduced in MCRS1-silenced cells with an average of 7.27 KMT (n=106, p-value=0.0002) instead of 9.19 KMTs per k-fiber (n=44) in siScramble spindles. No significant differences were found between this value in siScramble and control cells (8.93 KMTs per k-fiber in control cells, n= 226, p-value = 0.599). Altogether, the morphological alterations in MCRS1-silenced spindles support previously proposed changes in the dynamics of the MT minus ends (Meunier and Vernos, 2011).

**Figure 3.**
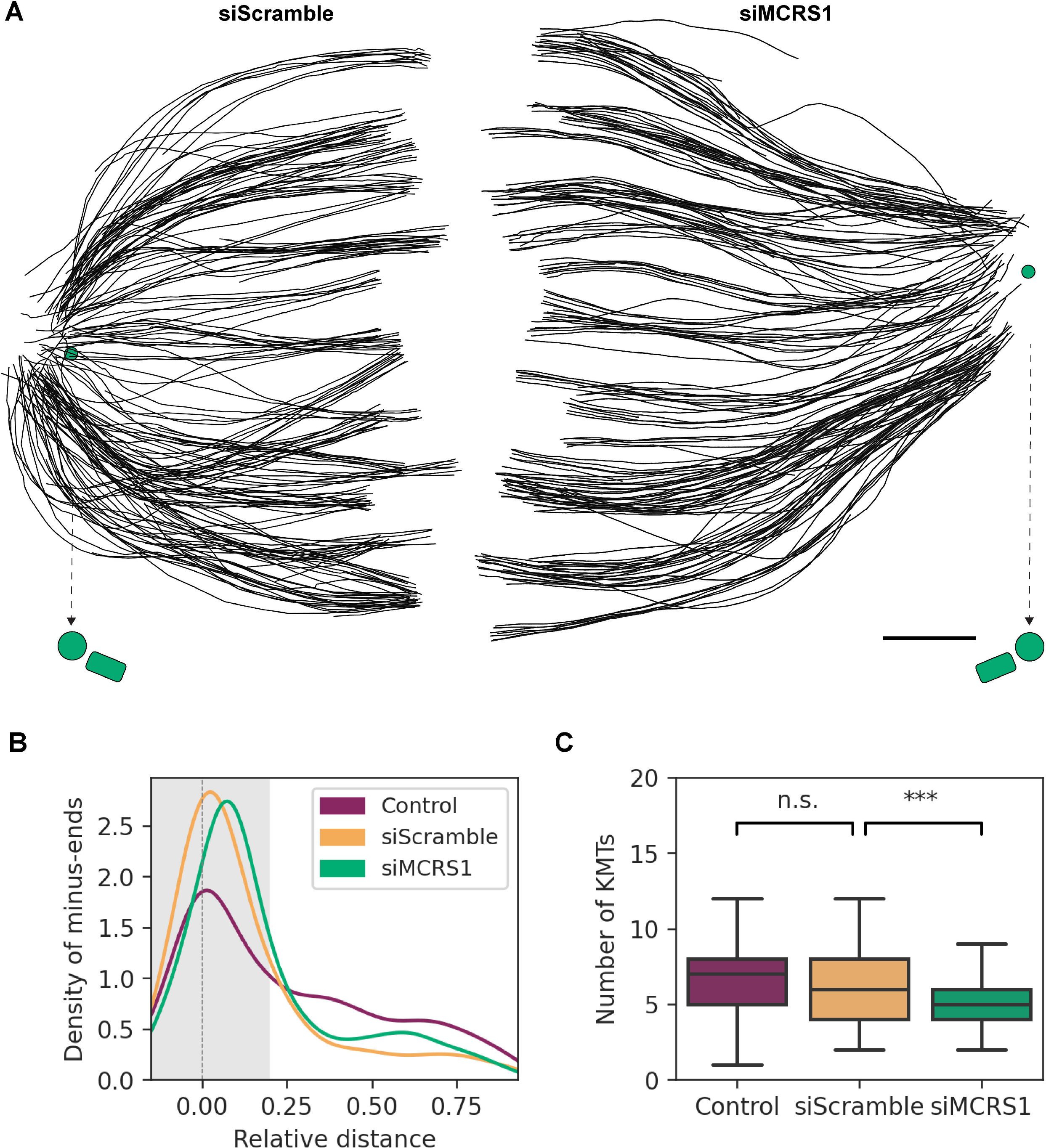
Shape of k-fibers in siScramble and MCRS1-silenced spindles. (**A**) Orthogonal projections (top views) of 3D reconstructions of quarter spindles in a siScramble (left, Scramble spindle #2) and a siMCRS1 cell (right, MCRS1 spindle #5) showing k-fibers as black lines. The spindle poles are indicated by green circles. The mother centrioles of the spindle poles are indicated by arrows (with dashed lines). Scale bar, 1μm. (**B**) Line-plot representing the Gaussian kernel density distribution of k-fiber minus ends at different relative distances from the mother centriole (position = 0) to the kinetochores (position = 1). The shaded area indicates the MT-centrosome association area. The shaded area represents the MT-spindle pole association area. (**C**) Boxplot representing the number of KMTs per k-fiber in the three conditions. The boxes show the upper and lower quartiles, the whiskers show the minimum and maximum values excluding outliers; the line inside the box indicates the median. Asterisks show significant differences according to a generalized linear model with a p-value = 2 × 10^−4^. N.s stands for non-significant with a p-value = 0.599 (Control: n=226, siScramble: n=44, siMCRS1: n=106).

### Silencing of MCRS1 increases the proportion of open minus ends of all spindle MTs

To explore more specifically the impact of MCRS1 silencing on the spindle MTs, we then quantified the morphology and relative distribution of their minus ends within the spindle. We selected the KMTs in the reconstructions of the siMCRS1 spindles as described above and manually classified their minus end morphologies into three categories: open, closed and undefined (**Fig. 1B; Fig. S5**). Next, we determined the relative distance of each KMT minus end along the half-spindle axis. The percentage of open KMT minus ends in the siMCRS1 spindles was 75.28% (CI = [44.83, 92.32]%, n=386) in the region surrounding the centriole pair (d < 0.2; **Fig.4A; Video 7**, green line), whereas it was 56.17% in control cells (CI = [25.05, 79.23]%, n=2071). To compare the two percentages, as percentages are bound between 0 and 100, we used the log2 transformation of the ratio open/closed. In this new scale, 0 means that there are the same number of open and closed minus ends; log2(open/closed) = 1 indicates that there are two times more open than closed minus ends, and −1 that there are two times more closed than open minus ends. This quantity is unbound and symmetric regardless of the group order, so it is more appropriate to represent differences between two groups in percentages. The log2(open/closed) was 1.38 higher in siMCRS1 cells compared to control cells around the centrioles (CI = [0.31, 2.39]; **Fig. 4B**). Our data show that MCRS1 reduction leads to an increase in the percentage of “uncapped” KMT minus ends.

**Figure 4.**
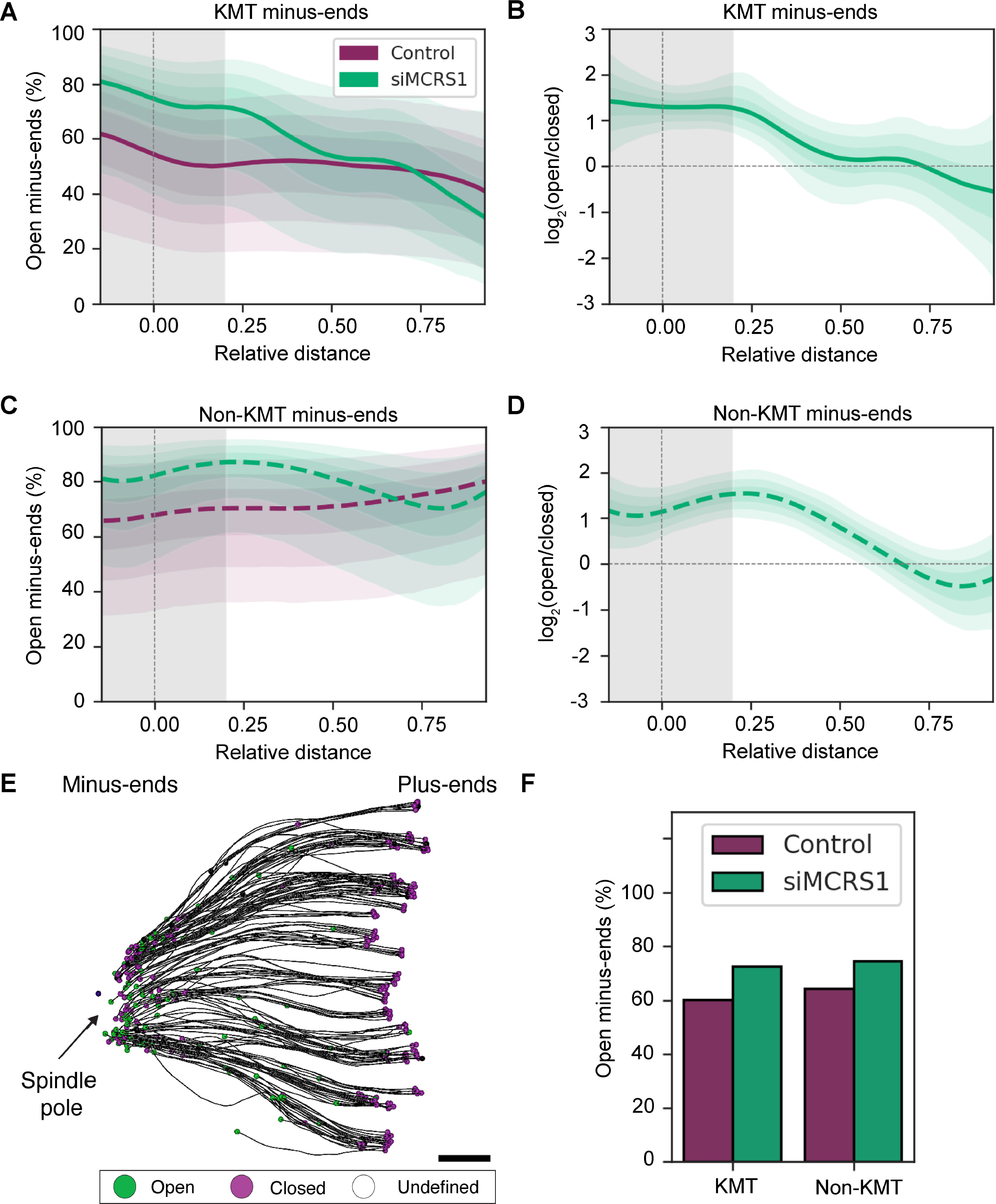
MT minus-end morphology in MCRS1-silenced cells. (**A**) Estimated percentage of open KMT minus ends plotted against the relative distance on the half-spindle length (position 0 = mother centriole, position 1 = kinetochores) in control (purple line; n=1707) and siMCRS1 cells (green line; n=541). The shadow around the lines (purple and green) represents the credible interval (CI) with a 95% probability to observe the true value within this interval. The shaded area (grey) represents the MT-spindle pole association area. (**B**) Posterior distribution of the comparison of the log2(open/closed) of siMCRS1 *versus* control (as shown in A) in KMTs (green line). The shadow around the line (green) represents the 95% credible interval (CI). (**C**) Estimated percentage of open non-KMT minus ends plotted against the relative distance on the half-spindle length in control (purple line, n=1084) and siMCRS1 cells (green line, n=3115). The shadow around the line (green) represents the 95% credible interval (CI). (**D**) Posterior distribution of the comparison of the log_2_(open/closed) of siMCRS1 *versus* control (as shown in C) in non-KMTs (green line). The shadow around the line (green) represents the 95% credible interval (CI). (**E**) Three-dimensional model (top view) showing KMTs in a quarter reconstruction of a siMCRS1 cell (siMCRS1 #5). KMTs are depicted as black lines and their end morphologies are indicated as open (green circles), closed (purple circles) or undefined (open circles). The position of the spindle pole is indicated by an arrow. Scale bar, 500 nm. (**F**) Bar representation of the estimated percentage of open KMTs (Control: 60.21%, n=2064; siMCRS1: 72.66%, n=848) and non-KMT minus ends (Control: 64.30%, siMCRS1: 74.51%, n=893; siMCRS1, n=4119) in control (purple) and siMCRS1 cells (green) in the spindle pole region (d< 0.2).

We then examined the minus end morphologies of non-KMTs at the spindle poles following the procedure described above. Due to the high number of this class of MTs, we performed this analysis on an unbiased randomly selected number of non-KMTs. We found that the estimated percentage of open non-KMT minus ends in the region near the centriole pair (d < 0.2) was 80.76% (CI = [53.16, 93.31]%) in MCRS1-silenced cells and 64.04% (CI = [33.00, 86.21]%) in control cells (**Fig. 4C**). As before, we used the log2 transformation of the ratio open/closed to compare both percentages. The log2(open/closed) was 1.35 higher in siMCRS1 cells compared to control cells for non-KMTs (CI = [0.56, 2.14], **Fig. 4D**) for the non-KMT minus ends. This analysis revealed that the non-KMT minus end morphologies are also altered close to the spindle pole in the absence of MCRS1.

Previous experiments suggested that MCRS1 function was specific for the regulation of the KMT minus end dynamics (Meunier and Vernos, 2011). To directly address this, we analyzed our data to determine whether the change in the ratios of close and open minus ends for KMTs and non-KMTs in control *versus* siMCRS1 cells was similar or not. We eliminated the distance-dependence of our analysis and restricted our quantification to the spindle-pole region (d < 0.2) for both MT populations (**Fig. 4F**). Considering both categories together, the change in log(open/closed) between siMCRS1 and control cells for non-KMTs was not significantly different from that in KMTs (Δlog(open/closed) = −0.07, p-value = 0.79). Altogether, these results suggested that MCRS1 silencing has a similar impact on the morphology of both KMT and non-KMT minus ends at the spindle poles with an increase in the percentage of open ends in the absence of MCRS1.

## Discussion

The depolymerization of MT minus ends has been postulated to play an essential role in the control of spindle size, k-fiber dynamics and chromosome movements (Barisic et al., 2021; Waters et al., 1996). However, there is currently little information on the precise mechanism that establishes and controls MT minus end dynamics at the spindle poles. An analysis of MT ends at spindle poles is hampered by the fact that the centrosomes are extremely dense locations in mitotic spindles, thus making it impossible to apply light microscopy for analysis of MT dynamics at the level of individual polymers. Direct visualization of MT ends by electron microscopy, however, can provide insightful clues about the end-morphology of individual MTs (O’Toole et al., 2003; Redemann et al., 2017). As an example, the majority of KMT plus ends are dynamic and show mainly open and flared morphologies (McIntosh et al., 2008; McIntosh et al., 2018; McIntosh et al., 2013; VandenBeldt et al., 2006), suggesting that there is indeed a correlation between MT end-morphologies and dynamic states (Hoog et al., 2011). In this context, we aimed to analyze MT minus end-morphology to infer their dynamics. For this, we applied large-scale electron tomography with single-MT resolution (Redemann et al. 2017; Kiewisz et al., 2022) and directly visualized MT minus end morphologies at mitotic spindle poles in human cells in metaphase.

Using this approach, we show that MT minus ends do not have homogeneous morphologies. We further show that both KMTs and non-KMTs display a mixture of both closed and open minus ends, in similar proportions (**Fig. 5**). The closed conformation that we observed for a large proportion of MT minus ends is reminiscent of previous reports on non-dynamic MT minus ends capped by the y-TuRC (Zheng et al., 1995). This suggests that the closed end morphology may correspond to stable and/or anchored MTs. The mixture of morphologies that we observed for the MT minus ends in the spindle may suggest that their depolymerization at the spindle pole is not synchronous, not even for KMTs within the same fiber.

**Figure 5.**
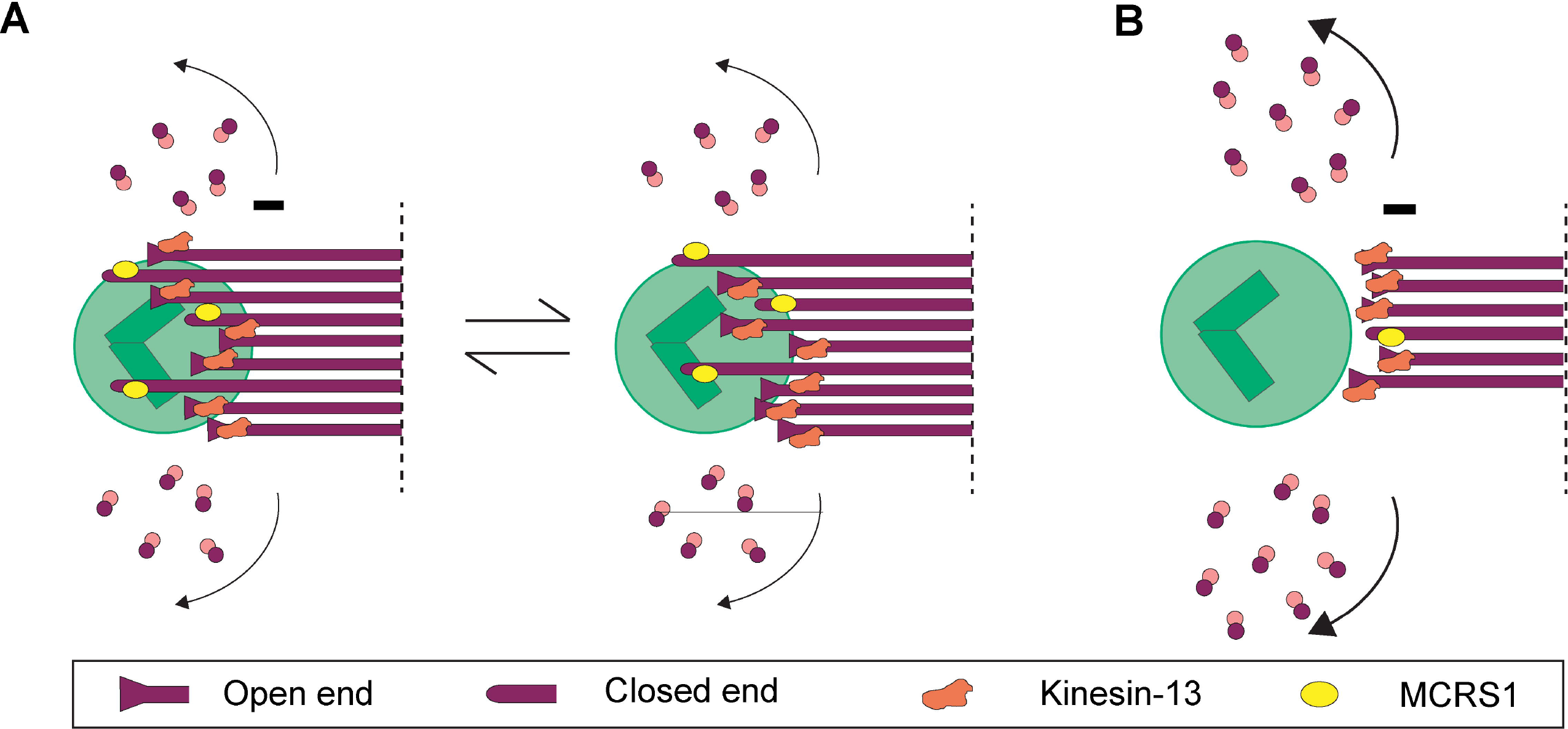
Proposed model of KMT minus-end dynamics regulation in mammalian metaphase. (**A**) Illustrates of the regulation of KMT minus-end dynamics in the metaphase of control cells. (**B**) Illustration of minus-end dynamics in response to MCRS1 silencing.

We observed a significant increase in the proportion of open minus ends at the spindle poles upon silencing of the proposed regulator of MT minus-end dynamics, MCRS1 (Meunier and Vernos, 2011). In addition, 3D reconstruction of spindles revealed an increase in the relative distance of the KMT minus ends to the centriole pair in these cells (**Fig. 4F**), which might be consistent with an increase in MT minus end depolymerization (Meunier and Vernos, 2011). Moreover, the loss of curvature of the KMTs (**Fig. 3A**, and **Fig. S7**) and the characteristic sigmoidal shape (recognized in all siMCRS1 spindles) suggest an increase in tension within the spindle, consistent with previous reports about the forces exerted by MT minus end depolymerization on the attached chromosomes (DeLuca et al., 2006; Waters et al., 1996).

MCRS1 was identified as a novel RanGTP-regulated factor that associates specifically with the k-fiber minus ends in cold-treated cells (Meunier and Vernos, 2011). We found here that both KMT and non-KMT minus-end morphologies change in MCRS1-silenced cells, suggesting that MCRS1 may in fact associate with both MT populations in the spindle. However, since it was recently reported that KMTs directly interact with many non-KMTs near the spindle pole (Kiewisz et al., 2022), it is possible that some of the MTs we classified as being non-KMT are in fact part of the k-fibers. Alternatively, MCRS1 may have a more general function in regulating MT minus end dynamics within the spindle than previously proposed.

It is interesting to speculate about the nature of the complex that associates with the MT minus ends to generate a cap structure. The γ-TuRC with an approximate molecular weight of 2353.97 kDa appears as a cone-like structure as observed *in vitro* (Brilot et al., 2021; Consolati et al., 2020; Drutovic et al., 2020; Keating and Borisy, 2000; Kollman et al., 2015; Moritz et al., 2000; Wieczorek et al., 2020; Wiese and Zheng, 2000; Zimmermann et al., 2020). The minus ends of some MTs may be capped by this complex and one would not expect these MTs to be dynamic. Instead, since MT minus ends are thought to depolymerize at the spindle pole, it is tempting to speculate that the electron-dense material that generates the close morphology could correspond to other MT minus-end binding complexes that could have more dynamic modes of binding, such as the MCRS1-KANSL-complex (Meunier et al., 2015), ASPM-katanin (Jiang et al. 2017), and/or NuMA/dynein (Elting et al., 2014) (**Fig. 5**).

Recent data suggested that MT minus end depolymerization and spindle flux can be uncoupled (Steblyanko et al., 2020). Instead, our data suggest that there is a direct correlation between the proportion of open MT minus ends and the speed of the spindle flux since it was reported to increase in absence of MCRS1 (Meunier and Vernos, 2011). The question remains, however, of how cells can undergo constant spindle flux, when MT minus ends are not synchronously depolymerizing. Unfortunately, our data can provide only a snapshot of the MT minus end morphologies in spindles at a given time point. Most likely, the binding/unbinding of components at the MT minus ends is stochastic. In such a model, MT minus ends would overall depolymerize, although individual MTs may pause, which would be visible as a heterogeneity in the morphology of individual ends (**Fig. 5**). A similar situation but with a net polymerization instead of a depolymerization has been described for MT plus ends (VandenBeldt et al., 2006). The putative mechanism underlying MT minus end dynamics at the spindle poles, therefore, still needs to be addressed in the future (Chen et al., 2019; Dudka et al., 2019).

## Experimental procedures

### Standard techniques

#### Cell culture

HeLa (Kyoto) cells were grown at 37°C in a 5% CO_2_ atmosphere. The cells were cultured in DMEM medium (Thermo Fisher Scientific, Waltham, MA) containing 4.5 g/L glucose and supplemented with L-glutamine with sodium pyruvate (Lonza, Basel, Switzerland), 10% fetal bovine serum (Thermo Fisher Scientific, Waltham, MA) and an antibiotic cocktail containing 100 μg/ml streptomycin and 100 units/ml penicillin (Sigma-Aldrich, St Louis, MO). For regular maintenance, HeLa cells were split with 0.25% Trypsin (Sigma-Aldrich, St Louis, MO) when reaching around 80-90% confluency.

#### RNA interference

Cells were seeded at 150.000 cells/ml in 75 cm^2^ flasks (Sigma-Aldrich, St Louis, MO) the day before transfection. Per flask, 500 pmol of siRNA and 25 μl Lipofectamine RNAiMax transfection reagent (Life Technologies, Carslbad, CA) were used and scaled accordingly for smaller volumes. Transfection was performed following the manufacturer’s protocol. After 48 hours post-transfection, cells were split and seeded. At 60 hours, cells were transfected as described above using siRNA purchased from Dharmacon (Lafayette, CO) as previously described (Meunier and Vernos, 2011) using the following sequences:

Scrambled: 5’-CGUACGCGGAAUACUUCGAUU-3’

MCRS1: 5’-GGCAUGAGCUCUCCGGAC-3’

Cells treated twice with siRNA as described above were collected after 72h either for Western Blotting and imaging (i.e: immunofluorescence microscopy or for electron tomography).

#### Gel electrophoresis and Western Blots

Protein lysates from HeLa cells were prepared by resuspending cell pellets in RIPA buffer containing 50mM Tris-HCl pH 7.4, 150 mM NaCl, 2 mM EGTA, 1% Triton X-100 (Sigma-Aldrich, St Louis, MO), 0.5% DOC (Sigma-Aldrich, St Louis, MO) and 0.1% SDS supplemented with protease inhibitors (Sigma-Aldrich, St Louis, MO). After incubation on ice for 15 minutes, the lysates were centrifuged at 4°C at maximum speed for 15 minutes. The protein content of the lysates was determined by using the Pierce BCA kit (Thermo Fisher Scientific) and following the manufacturer’s instructions. For each run, 50 μg of protein lysate was diluted in loading buffer 5x, boiled for 10 minutes and loaded in a 10% SDS-PAGE gel. Gels were run at 120 V for 90 minutes.

For Western Blots, a semi-dry transfer was done to blot the proteins in a 0.45 μm nitrocellulose membrane (GE Healthcare, Chicago, IL). Proteins were transferred for 90 minutes at 65 mA. The membrane was blocked using 5% milk (Sigma-Aldrich, St Louis, MO) in TBS at 4°C overnight. Primary antibodies (MCRS1, home-made; Tubulin (DM1A), T6199, Sigma-Aldrich, St Louis, MO) were incubated in 0.1% Tween20 (Sigma-Aldrich, St Louis, MO) in TBS for 1 hour at room temperature. Secondary antibodies conjugated with AlexaFluors (AlexaFluors800 – MCRS1; AlexaFluors680 - α-Tubulin) were incubated in the same buffer at room temperature for 45 minutes. Blots were developed using an Odyssey Infrarred Imaging System (Li-cor, Lincoln, NE).

### Light microscopy

For immunofluorescence, HeLa cells were grown on 18-mm round coverslips (Marienfeld, Germany) in six-well plates. Cells were fixed by immersion in cold methanol (−20°C) for 10 minutes. Cells were then blocked and permeabilized in 0.5% BSA (Panreac, Barcelona, Spain) and 0.1% Triton X-100 (Sigma-Aldrich, St Louis, MO) in 1x PBS for 30 minutes at room temperature. Primary antibodies (MCRS1, homemade; Tubulin (DM1A), T6199, Sigma-Aldrich, St Louis, MO) were incubated in the same blocking buffer for 1 hour at room temperature. Secondary antibodies conjugated with AlexaFluors 488 and Hoechst to stain DNA (Life Technologies, Carlsbad, CA) were diluted in blocking buffer 1:1000 and incubated at room temperature for 45 minutes. Coverslips were mounted. in 9.6% (w/v) Mowiol, containing 24% (w/v) glycerol (Sigma-Aldrich, St Louis, MO) and 0.1 M Tris-Cl).

After immunostaining, samples were visualized by light microscopy to select MCRS1-depleted cells. The samples were visualized with a LEICA SP5 microscope (Leica Microsystems, Austria) equipped with a water 63x objective lens (Leica HCX PL APO 63x/1.2 W) and a PMT Leica hybrid detector camera (HyD) (Dresden). Images were acquired using a 488 nm argon laser with a HyD sensor set to collect light from 500nm to 550 nm and a 594 nm diode laser with a sensor set from 620 nm to 730 nm. Chromosome alignment at the metaphase plate was considered as an indicator of the mitotic stage. Selected mitotic cells in the metaphase stage were further characterized by measuring both the half-spindle and the MT angle using Fiji (Schindelin et al. 2012) (**Figure 2B**). For this, the maximum intensity projection tool was used for the z-stacks corresponding to the α-tubulin staining in the selected spindles. A threshold was set for the projected images to create a binary mask. The angles were measured manually using the angle tool. The half-spindle angle was measured to analyze the shape of the spindle poles, and it was measured by calculating the angle between the metaphase plate and the spindle pole (**Figure 2B**). The MT angle was measured to analyze the degree of bending of the outer MTs. We measured the outline of the quarter-spindle by calculating the angle between the metaphase plate and the spindle pole (**Figure 2B**). Differences in the measured angles in both siScramble and siMCRS1 spindles were assessed by applying a linear regression model.

### Electron microscopy

#### Sample preparation

In preparation for electron microscopy, 3-mm and 6-mm sapphire discs (M. Wohlwend GmbH, Switzerland) were prepared for the attachment of cells. The mitotic fraction of the cell culture was then collected by applying the “shake-off” technique (Kiewisz, Müller-Reichert, and Fabig 2020). Briefly, flasks with HeLa cells were subjected to two “shake-off” rounds. Flasks were hit against the surface of the bench to detach mitotic cells. The collected cells were then centrifuged at 1200 rpm at 37°C for 4 minutes and the cell pellets were resuspended in 1 ml of DMEM medium (Thermo Fisher Scientific, Waltham, MA) supplemented with 10% bovine serum albumin (BSA; Thermo Fisher Scientific, Waltham, MA) and 10% of fetal bovine serum (Thermo Fisher Scientific, Waltham, MA). Next, the cells were allowed to attach to carefully cleaned sapphire discs for 10 minutes at 37°C. Cleaning of the sapphire discs included immersion in a Piranha solution (1:1 H2SO4 and H202, v/v), coating with poly-L-lysine 0.1% (w/v) and drying at 60°C for 2 hours. Finally, the discs were incubated with a 1:10 solution of fibronectin (Sigma-Aldrich, St Louis, MO) in PBS at 37°C for 2 hours prior to use.

#### High-pressure freezing and freeze substitution

Cells attached to 3-mm sapphire discs were cryo-immobilized using an EM ICE (Leica Microsystems, Austria) and 6-mm sapphire discs were frozen by using a Compact 03 high-pressure freezer (M. Wohlwend GmbH, Switzerland). For each freezing round with the EM ICE, a type-A aluminum carrier (Leica Microsystems, Austria) with the 100 μm-indentation facing up was placed in the specimen loading device of the freezer. The cavity of the carrier was then filled with 5 μl of DMEM containing 10% BSA and the sapphire disc with the attached HeLa cells facing down was placed onto the carrier. Next, a spacer ring was mounted on top of the sample and the freezing started immediately. For using the Compact 03 high-pressure freezer, the 6-mm sapphire discs with the mitotic cells facing down were placed on a type-A aluminum planchette with the 40μm-deep cavity pre-filled with warm DMEM medium supplemented with 10% BSA. Closed carriers were then placed in the specimen holder, clamped with the holder arm and immediately cryo-immobilized. This approach as developed for freezing with the Compact 03 allowed for a quick inspection of the assembled specimens, so that only samples without trapped air were further processed for a high-pressure freezing (Kiewisz et al., 2022). Using both freezers, samples were frozen at ~2,000 bar with a cooling rate of ~20,000°C/s (Reipert, Fischer, and Wiche 2004). After freezing, samples were stored in liquid nitrogen until further use. Freeze substitution was carried out as previously described (Muller-Reichert et al., 2003). Briefly, samples were transferred to cryo-vials filled with anhydrous acetone containing 1% osmium tetroxide (EMS, USA) and 0.1% uranyl acetate (Polysciences, USA). Freeze-substitution was done using an automatic freeze substitution machine (EM AFS, Leica Microsystems, Germany). Samples were kept at −90°C for 1 hour, then warmed up to −30°C in steps of 5°C per hour and maintained at −30°C for 5 hours. Next, the temperature was increased to 0°C in steps of 5°C per hour. Finally, after reaching 0°C samples were washed three times with pure anhydrous acetone at room temperature.

#### Sample embedding, pre-selection of mitotic cells and ultramicrotomy

For resin embedding, the samples were placed in flow-through chambers (Leica, Germany) and infiltrated with Epon/Araldite (EMS, Hatfield, PA) in three steps of 1 hour each with increasing concentration of resin: 1:3, 1:1 and 3:1 (resin:acetone, w/v), followed by a single step overnight in pure resin at room temperature (Müller-Reichert et al. 2003). Then, samples were polymerized at 60°C for 48 hours. After polymerization the plastic samples were removed from the flow-through chambers. Using a razor blade, the sapphire discs were then removed from the resin blocks to expose the embedded cells for further processing as described previously (Kiewisz et al., 2022). This procedure was chosen to avoid any re-mounting of thin layers of resin on dummy blocks.

In order to screen for cells in metaphase, the resin blocks were observed from the top using an upright brightfield microscope (Zeiss, Germany). The two main features used to select cells in metaphase were a rounded shape and a distinguishable metaphase plate (Kiewisz et al., 2021). Serial semi-thick (300 nm) sections were cut using an EM UC6 ultramicrotome (Leica Microsystems, Austria) and collected on Formvar-coated slot grids. Samples were post-stained with 2% uranyl acetate for 10 minutes followed by a 0.4% of Reynold’s lead citrate solution (Science Services, Germany). Colloidal 15nm-gold particles (British Biocell International, UK) were attached to both sides of the sections mounted on the grids to serve as fiducial markers for tomographic reconstruction.

#### Final staging of pre-selected spindles

The serial sections were imaged using a TECNAI T12 Biotwin transmission electron microscope (ThermoFisher Scientific, Waltham, MA) operated at 120kV and equipped with an F214 CCD camera (TVIPS GmbH, Germany). Images of whole cells in metaphase were acquired at 1200x magnification using the EMMenu Software (TVIPS GmbH, Germany). To choose a cell for electron tomography, the metaphase plate had to be correctly formed when looking at the chromosome area in 3D, and for each chosen cell the half spindle and the MT angle were measured in the 3D volumes as described (see the section on immunofluorescence). For this, the EM stacks were projected in 3D. The chromosome and microtubule area were estimated by manually labeling the chromosome and MT area. Both the half-spindle angle and MT angle were finally calculated using the ZIB Amira software (Zuse Institute Berlin, Germany).

### Three-dimensional reconstruction by electron tomography

#### Data acquisition and calculation of tomograms

Electron tomography was performed on the selected metaphase cells as previously described (Kiewisz et al. 2021). Briefly, a series of tilted views were recorded using a TECNAI F30 transmission electron microscope (ThermoFisher Scientific, Waltham, MA) operated at 300 kV and equipped with a Gatan US1000 2K × 2K CCD camera. The SerialEM software package was used for the acquisition and montaging of data sets (Mastronarde, 2003; Mastronarde, 2005). For dual-axis electron tomography, images were captured every 1.0° over a ±60° range at a pixel size of 2.32 nm. For a recording of the second axis, grids were rotated by 90° and another series of tilted views was acquired (Mastronarde, 1997). Tomograms were calculated by using the IMOD software package (Mastronarde and Held, 2017). To increase the sample number, we acquired volumes of quarters of the cells (see **Table 1**).

#### Segmentation of MTs and stitching of serial tomograms

MTs were semi-automatically segmented using the ZIB Amira (Zuse Institute Berlin, Germany) software package (Lindow et al., 2021) as previously described (Lindow et al., 2021; Redemann et al., 2014; Weber et al., 2012). After manual correction of MT segmentation, the serial tomograms of each recorded cell were stitched using the segmented MTs as alignment markers (Lindow et al., 2021; Weber et al., 2014).

#### Z-correction of stacked tomograms

Each stack of serial tomograms was expanded in *Z* to correct for a sample collapse during the data acquisition (McEwen and Marko, 1999). We corrected this shrinkage by applying a Z-factor to the stacked tomograms (Kiewisz et al., 2022; Kiewisz et al., 2021; O’Toole et al., 2020). Taking the microtome setting of around 300 nm, we multiplied this value by the number of serial sections. For each spindle, we also determined the thickness of each serial tomogram and then calculated the total thickness of the reconstruction. The *Z*-factor was then determined by dividing the actual thickness of each stack of tomograms by the total thickness as determined by the microtome setting. Such calculated Z-factors were then applied to our full spindle reconstructions.

### Quantitative analysis of tomographic data

For quantitative analysis of the tomographic data, the automatic spatial graph analysis (ASGA) software tool (Kiewisz and Müller-Reichert, 2022) was used to measure the outer-kinetochore distance, the number of MTs per k-fiber, and the distribution and morphology of the MT minus ends. Quantitative analyses were carried out essentially as described (Kiewisz et al., 2022).

#### Outer-kinetochore distance

The inter-kinetochore distance was used as a readout of the mitotic stage of the selected cells (Maresca and Salmon, 2009). To measure the inter-kinetochore distance, the neighboring sister kinetochores were identified in the 3D models. The center of each kinetochore was defined as the median position of all KMT plus ends associated with each selected outer kinetochore, and the inter-kinetochore distance was then calculated as the 3D Euclidean distance between the defined median centers of each kinetochore pair. The inter-kinetochore distances were determined for control, siScramble and siMCRS1 cells, and values were compared using a linear regression model.

#### Number of KMTs per k-fiber

MTs associated with the kinetochores were defined as KMTs. Accordingly, the average number of KMTs was determined for each experimental condition. Analysis of the number of KMTs per k-fiber was done using a Generalized Linear Model (GLM) in R, applying a Poisson likelihood and a log link function. The Poisson distribution is generally used to model discrete positive outcomes such as the number of microtubules when the numbers are small and cannot be approximated by a normal distribution (n < 20).

#### Distribution of MT minus ends

The polarity of KMTs was assigned as follows. The end of each KMT associated with a kinetochore was assumed to be the plus end and the other end the minus end. To analyze the position of the KMT minus ends in the metaphase spindles, the 3D Euclidean distance of each KMT end to both spindle poles (i.e., to the center of the mother centriole of the respective pole) was determined. Then, the KMT minus end was determined as the end closest to one of the spindle poles. In addition, the relative position of KMT minus ends along the pole-to-pole axis was calculated. The relative position of each minus end is given as the normalized position between the kinetochore (position = 1) and the mother centriole (position = 0) along the pole-to-pole axis. The distribution of relative positions of KMT minus ends (mean, ±SD) is as an average density distribution for each condition. For non-KMTs, the end closer to the nearest centriole pair was defined as the minus end. The relative distance for the non-KMTs was calculated between the average kinetochore (position = 1) and the mother centriole (position = 0).

#### Tortuosity of KMT

To analyze the global tortuosity of the KMTs, the ratio of the spline length and the 3D distance between the plus and the minus end of each KMT was measured (Kiewisz et al., 2022). The correlation of KMT tortuosity and their length is shown by a fitted curved line calculated with a logarithmic function.

#### Morphology of MT ends

MT ends were annotated by manual segmentation. MT ends were manually classified as open, closed or undefined by two different observers using the end classifier in the ZIB Amira software package (Detlev et al., 2005). To ensure an unbiased manual annotation of the end morphology, a random 3D view of each MT was presented to the analyst without knowledge of the MT identity (either KMT or non-KMT). The polarity of the MT ends (either plus or minus) and location (relative distance) within the spindle were determined after the classification of the end morphology.

Analysis of the MT end morphology was modeled as a binary outcome (open *vs.* closed), such that the number of open ends was naturally drawn from a binomial distribution depending on the true unobserved proportion of open ends. To model the dependency of the proportion of open ends on the distance to the spindle pole, we discretized the data into 16 different intervals and counted the number of open ends called by each observer. We used a third-order b-splines on the underlying logit transformation of the proportion of open ends to jointly infer the proportion of open ends at each possible distance from the pole. We then added the observer as a random effect to consider the differences in the classification made by the two independent observers.

#### Error analysis of tomographic data

Errors in automatic MT segmentation and in the stitching of serial tomograms have been discussed previously (Kiewisz et al., 2022; Kiewisz et al., 2021; Lindow et al., 2021; Redemann et al., 2014). As for the segmentation of MTs, the error associated for our approach is in the range of 5-10% (Weber et al. 2012). Each individual MT in our reconstruction has been checked manually for the correct tracing of both ends.

In previous publications (Lindow et al., 2021; Redemann et al., 2017; Weber et al., 2014), we estimated the overall quality of the MT stitching by analyzing the distribution of MT endpoints in the Z-direction (i.e., normal to the plane of the slice). We expect to find approximately the same density of MT endpoints along the Z-direction of each serial-section tomogram. Therefore, if the density of endpoints after matching is approximately the same along the Z-direction of the serial-section tomograms, we can assume that the number of artificial points that have been introduced at the interfaces of the serial sections are negligible (Kiewisz et al., 2022; Kiewisz et al., 2021).

## Data availability

Tomographic data has been uploaded to the TU Dresden Open Access Repository and Archive system (OpARA) and is available as open access: http://doi.org/10.25532/OPARA-XX

The code used to perform quantitative analysis of MT organization in spindles has been uploaded to the GitHub repository and is available as open access under the GPL v3.0 license: https://github.com/RRobert92/ASGA; https://github.com/RRobert92/ASGA_3DViewer

Data and code for statistical analysis of MT ends can be accessed at https://bitbucket.org/cmartiga/kfibers/src/master/

## Acknowledgments

The authors would like to thank support form the European Union’s Horizon 2020 research and innovation programme under the Marie Skłodowska-Curie grant agreement No 675737 (DiviDE ITN network) to A.L.D., R.K, I.V and T.M.-R. Research in the Müller-Reichert laboratory is supported by funds from the Deutsche Forschungsgemeinschaft (MU 1423/8-2). Work in the Vernos lab was supported by the Spanish Ministry of Economy (MINECO) I+D grant BFU2012-37163 and BFU2015-68726-P. A.L.D also received an EMBO short-term fellowship to visit the Müller-Reichert lab, grant agreement No. 8704. We thank Dr. Tobias Fürstenhaupt (Electron Microscopy Facility at the MPI-CBG, Dresden, Germany) for technical support. We also thank the Vernos and Müller-Reichert groups and the members of the DiviDE ITN for discussions. We acknowledge the Spanish Ministry of Economy, Industry and Competitiveness (MEIC) to the EMBL partnership and support of the Spanish Ministry of Economy and Competitiveness, ‘Centro de Excelencia Severo Ochoa’ as well as support of the CERCA Programme / Generalitat de Catalunya.

## Supplementary figures

**Figure S1.**
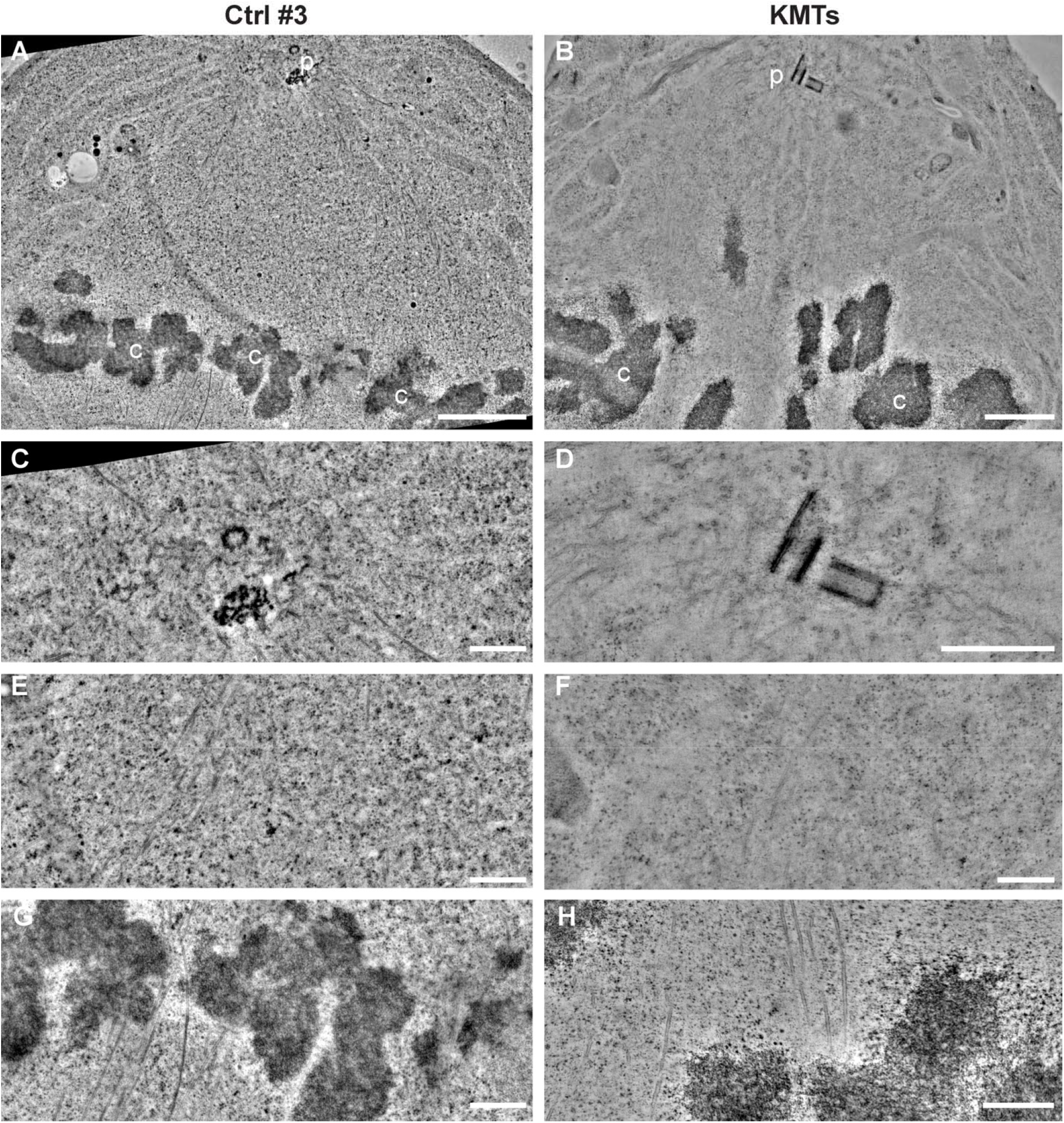
Illustration of metaphase in control and siMCRS1 cells. Tomographic slices are shown to illustrate the morphology of control (Ctrl #3, left column) and siMCRS1 cells (#5, right column). (**A–B**) Low-magnification images showing regions between a spindle pole (p) to chromosome (c) region. Scale bar, 1 μm. (**C–D**) Centrioles and MTs at higher magnification. Scale bars, 500 nm. (**E–F**) Regions between the spindle poles and the chromosomes. Scale bars, 500 nm. (**G–H**) Kinetochores with associated KMTs. Scale bars, 500 nm.

**Figure S2.**
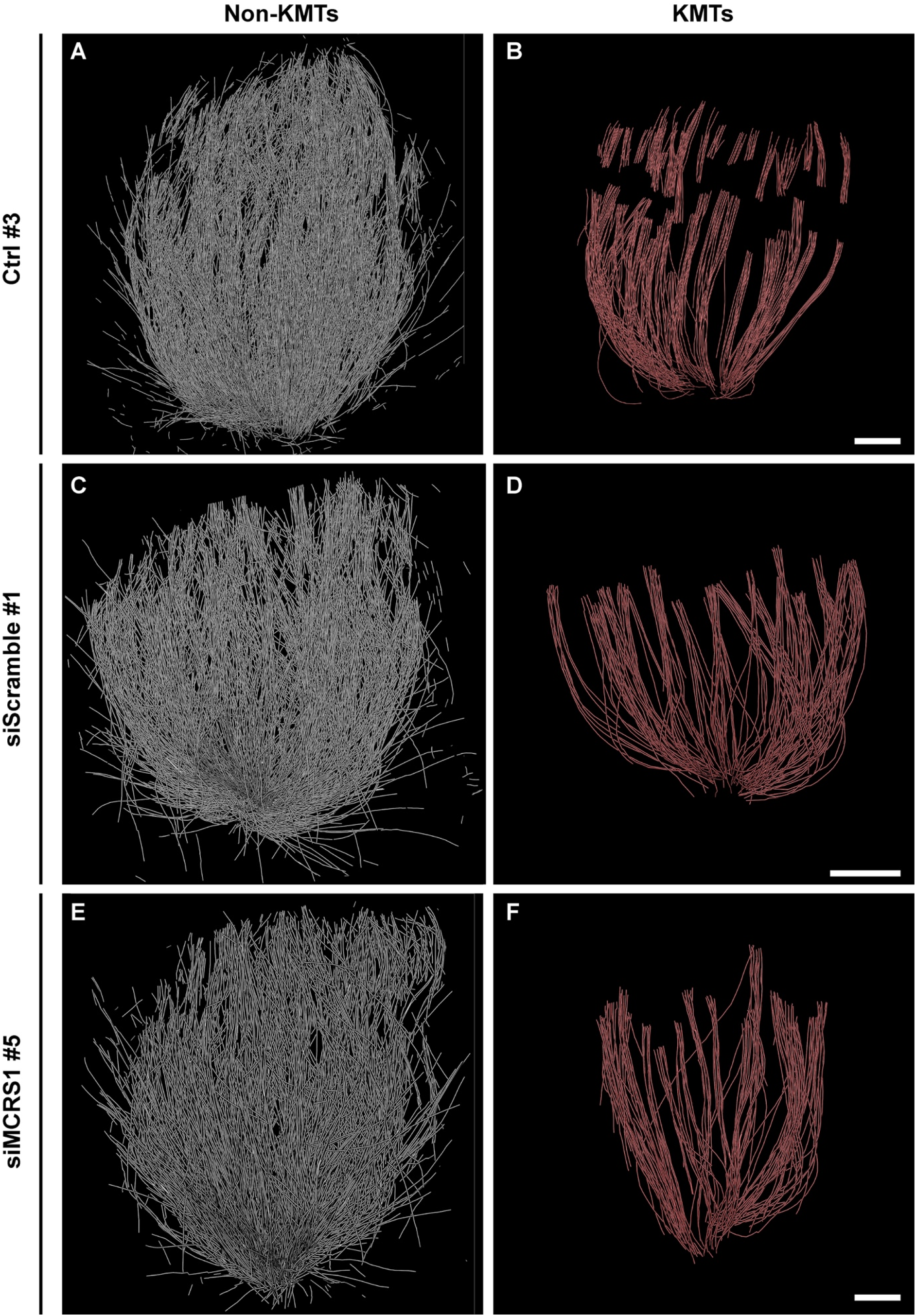
Three-dimensional modes of metaphase spindles as observed by large-scale electron tomography. (**A**) Three-dimensional model of non-KMTs in a control spindle (#3). (**B**) Illustration of only KMTs in a control spindle (#3) as shown in B. (**C**) Model of non-KMTs in a Scrambled spindle (#1). (**D**) Illustration of only KMTs in the Scrambled spindle (#1) as shown in C. (**E**) The model of non-KMTs in a siMCRS1 spindle (#5). (**F**) Illustration of only KMTs in the siMCRS1 spindle (#5) as shown in D. Scale bars, 1 μm.

**Figure S3.**
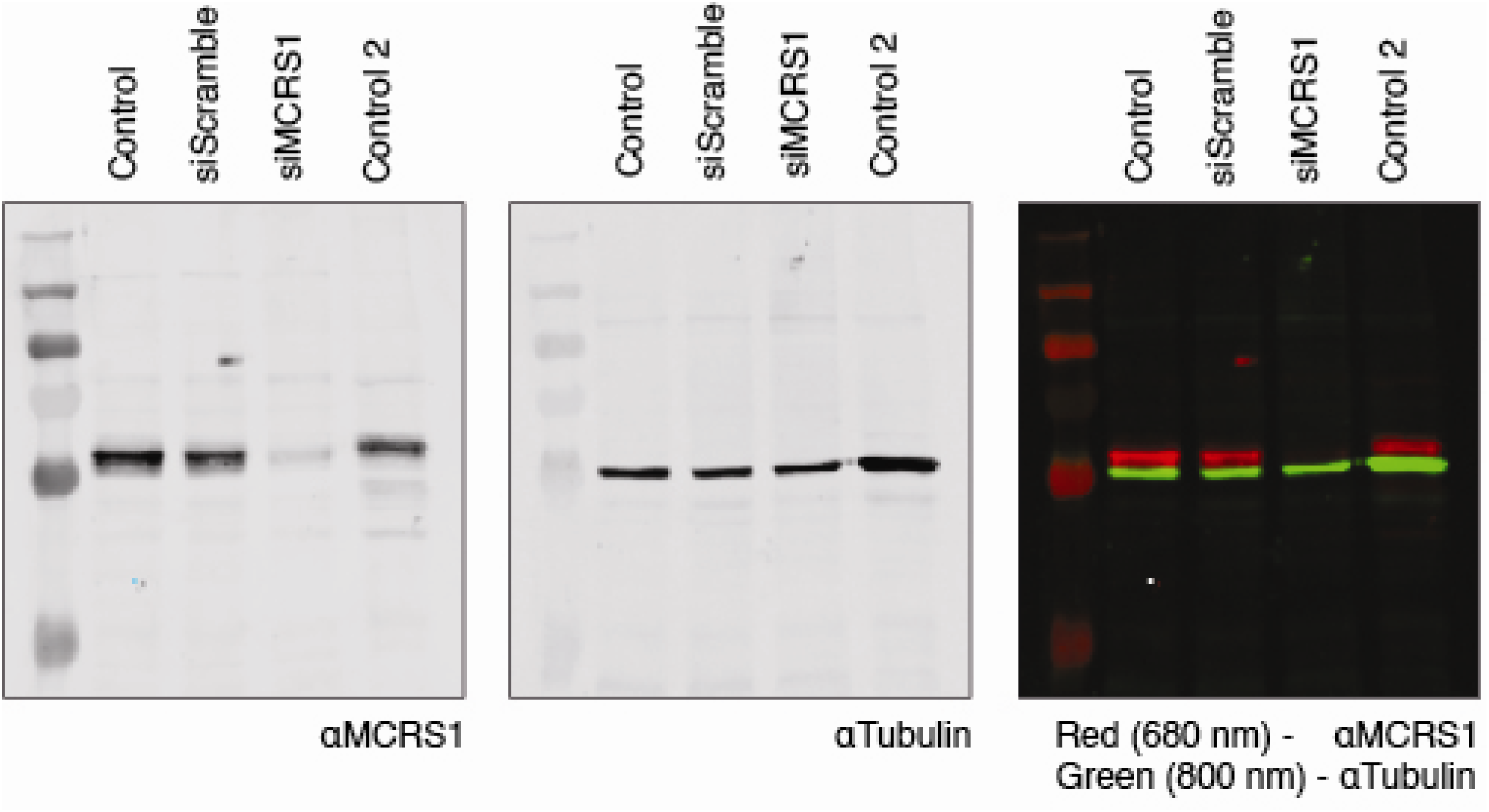
Analysis of protein levels by Western Blot. Full western blots corresponding to the cropped Western Blot shown in **Fig 1A**. Lysates from two controls, siScramble and siMCRS1 cells were run on SDS-PAGE. Left, the membrane was probed with the anti-MCRS1 antibody and re-probed with an anti-αTubulin antibody as loading control (middle). The overlay of both signals is shown on the right (anti-MCRS1, red; anti-Tubulin, green).

**Figure S4.**
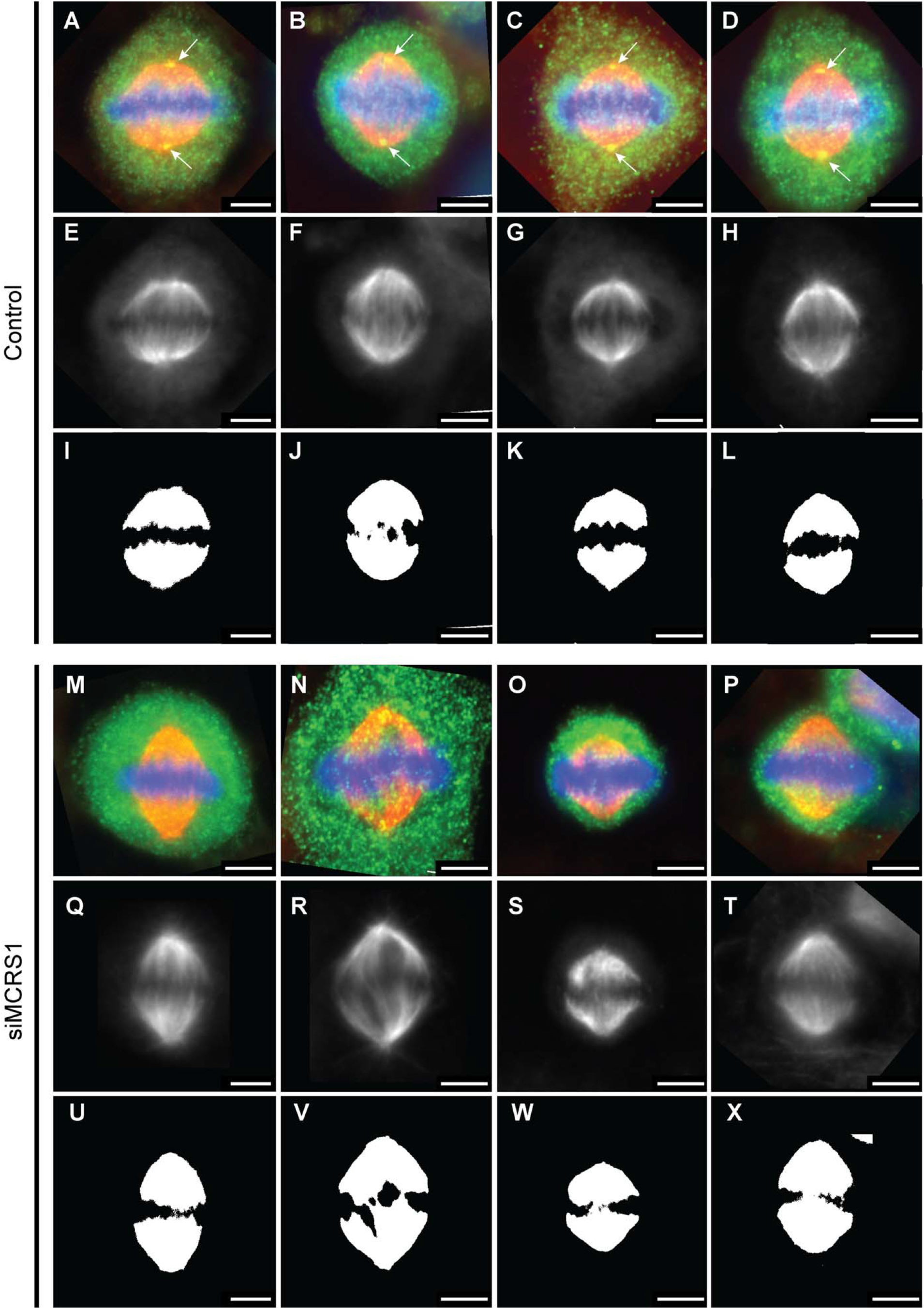
Angle analysis for control and siMCRS1 depleted HeLa cells. Set of fluorescent images showing the steps analysis. (**A-D**, and **M-P**) Raw images. MCRS1 is shown in green, DNA in blue and tubulin in magenta. MCRS1 staining can be observed as tiny spots at spindle poles in the control cell (arrows) (**E-H**, and **Q-T**) The middle row shows extracted tubulin channel after z-projection. (**I-L**, and **U-X**) Automatic threshold image. A created binary mask is used to define spindle and MT angles. Scale bars, 5 μm.

**Figure S5.**
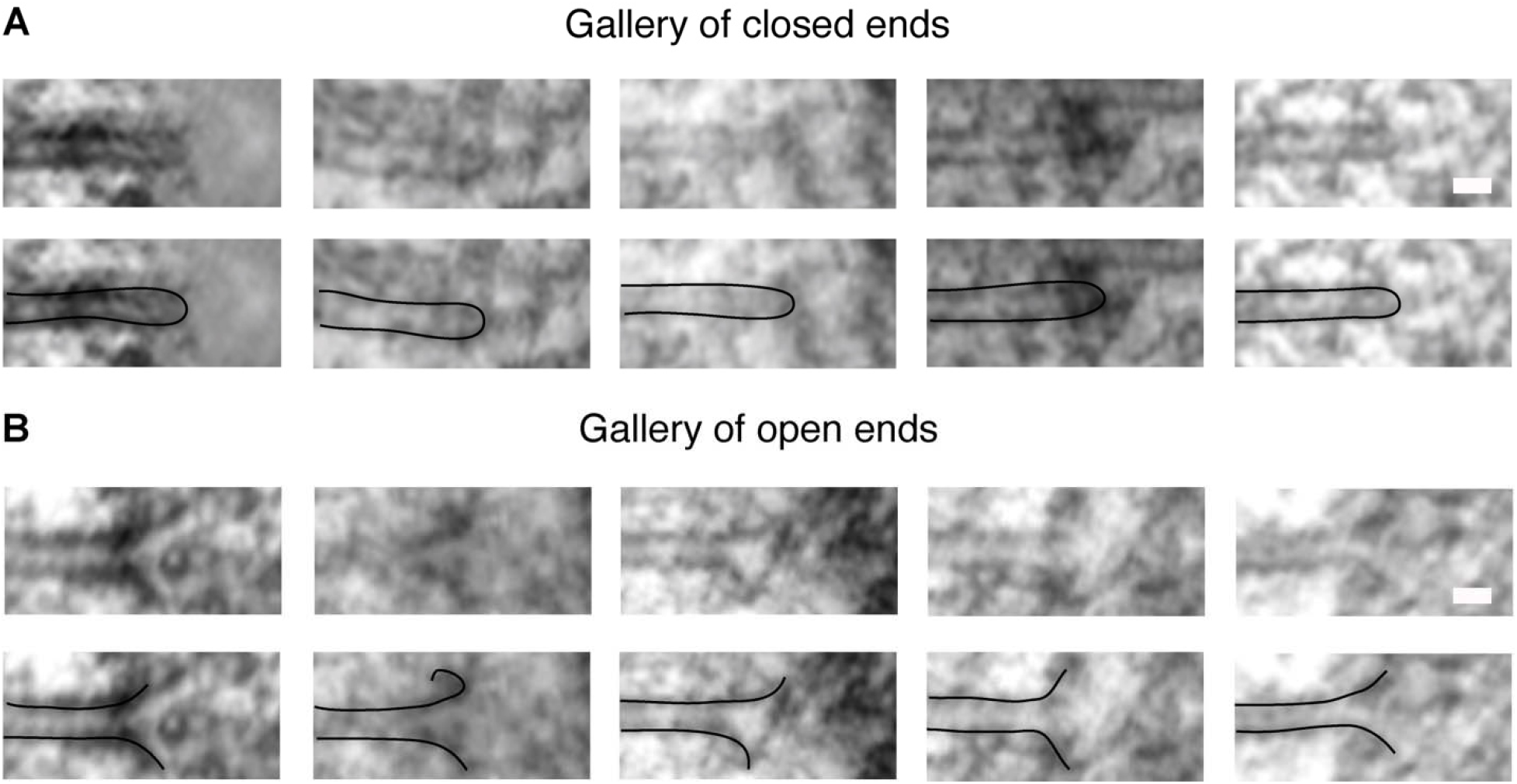
Gallery of MT ends morphology as observed by electron tomography. (**A**) Representative collection of images showing closed MT ends (top row, tomographic slices; lower row, tomographic slices with overlays). The polarity of the MTs is not assigned. Scale bar, 25 μm. (**B**) Collection of images showing open MT ends. Scale bars, 25 μm.

**Figure S6.**
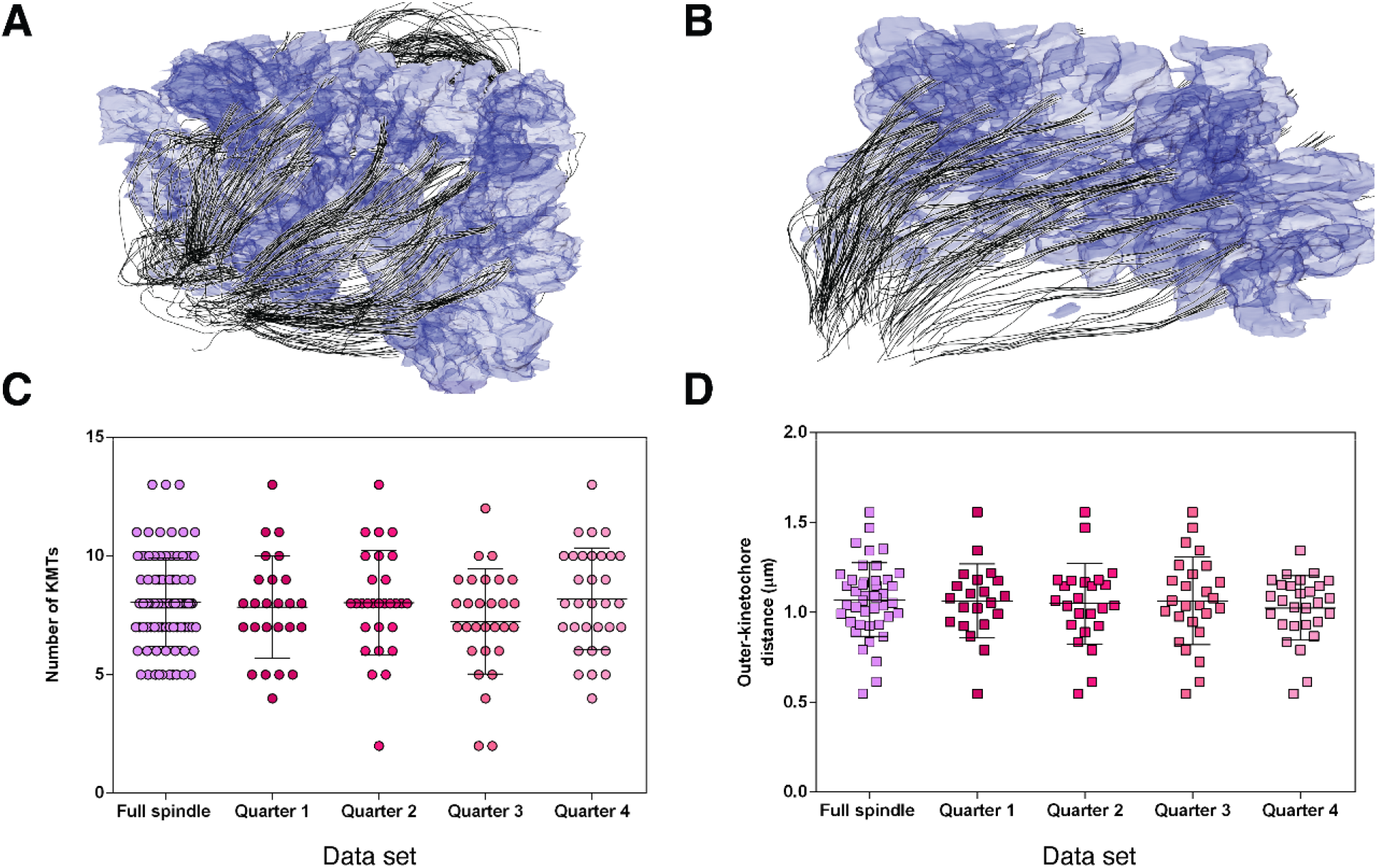
Comparison of the characteristics of a full spindle *versus* their quarters. (**A**) Perspective view of a full 3D model showing the KMTs in a control spindle #1. KMTs are shown as black lines, with chromosomes in gray. (**B**) Perspective view of a 3D model showing the KMTs in a quarter spindle #1. (**C**) Scatterplots showing the number of KMTs per k-fiber for a full spindle #1 and the four-quarters of the same spindle. The midline shows the mean, and the error bars represent ±SD. The mean numbers of KMTs for the full spindle is 8.04 ±1.86 μm (mean ±SD, n=93), and for the quarter (#1: 7.84 ±2.15 μm, n=25; #2: 8.03 ±2.20 μm, n=29; #3: 7.24 ±2.21 μm, n=29; and #4: 8.19 ±2.15 μm, n=32). There are no significant differences among the mean values for the full spindle and the quarter spindles after ANOVA analysis (p-value = 0.3791). (**D**) Scatterplot of the outer-kinetochore distance in the full spindle *versus* the spindle quarters. The midline shows the mean, and the error bars represent ±SD. The mean outer-kinetochore distances for the full spindle is 1.07 ±0.20 μm (n=43), and for the quarter (#1: 1.06 ±0.20 μm, n=21; #2: 1.05 ±0.23 μm, n=24; #3: 1.06 ±0.24 μm, n=26; and #4: 1.03 ±0.18 μm, n = 28). There are no significant differences among the mean values for the full spindle and the quarter spindles after ANOVA analysis (p-value = 0.9192).

**Figure S7.**
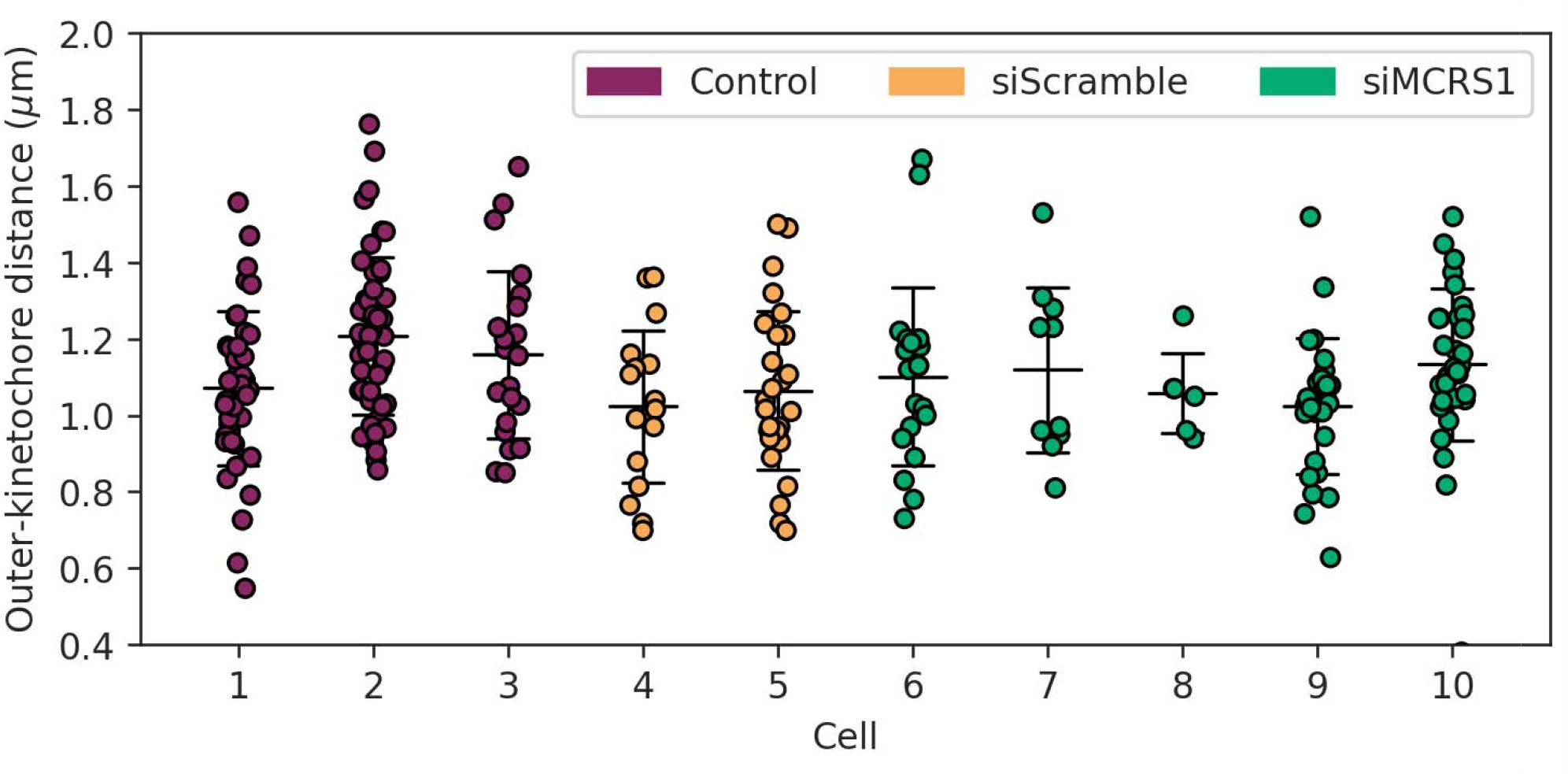
Comparative analysis of outer-kinetochore distances in 3D models of control, siScramble and siMCRS1-depleted cells. Scatter plots of the inter-kinetochore distances as measured for each of the 3D-reconstructed spindles. The mean inter-kinetochore distance is 1.14 μm (n=114) in control cells, 1.05 μm (n=44) in siScramble cells and 0.99 μm (n=98) in siMCRS1 cells. A linear regression model did not reveal significant differences between siScramble versus Control p-value = 0.108 and siScramble versus siMCRS1 (p-value=0.240). The numbers for each cell are matching the data sets as given in Table 1. Cells #1-2 correspond to full spindle reconstructions and cells #3-10 to quarter spindles.

**Figure S8.**
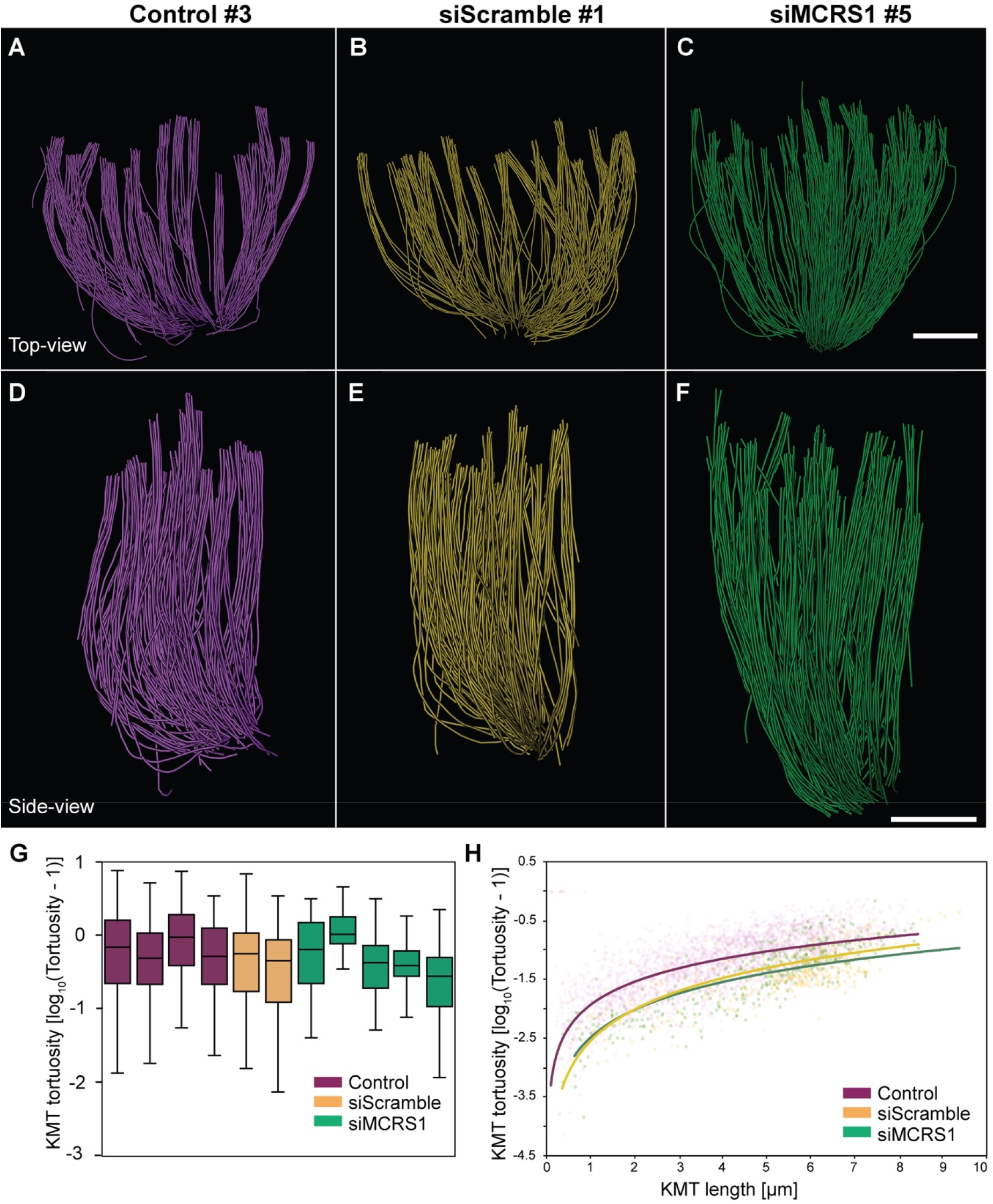
Analysis of KMT tortuosity in a 3D model of control, siScramble and siMCRS1 cells. (**A-C**) Orthogonal projections (top views) of KMTs in representative 3D models of ctrl (#3, left panel, lines in magenta), siScramble (#2, mid panel, lines in yellow) and siMCRS1 (#5, right panel, lines in green). Scale bar, 1 μm. (**D-F**) Side-views of the orthogonal projections of KMTs as shown in A-C. Scale bars, 1 μm. (**G**) Bar plot showing the logarithmic representation of KMT tortuosity as measured for each of the spindles. The boxes show the upper and lower quartiles, the whiskers show the minimum and maximal values excluding outliers; the line inside the box indicates the median (Control: n=3164, siScramble: n=408, siMCRS1: n=727). (**H**) Line plot showing a logarithmic representation of KMT tortuosity against the KMT length for each data point. The lines show a logarithmic tradeline for each condition.

## Supplementary videos

**Video 1. Generation of a 3D model assembled by joining serial electron tomograms**

Series of tomograms and corresponding 3D model of a control spindle #3. The video illustrates the stacking of serial tomograms to increase the volume of reconstruction. KMTs are shown as red lines. This video corresponds to the spindle shown in **Fig. 1C**.

**Video 2. Three-dimensional model of a low-resolution spindle as used for the analysis of spindle geometry in a siScramble cell**

Series of electron micrographs of a spindle in a siScramble cell #2. The stacking of serial tomograms to generate a 3D model of the spindle volume is illustrated. The spindle on each section is indicated by a transparent gray area. Chromosomes are shown in blue. This video corresponds to the spindle shown in **Fig. 2D, left panel**.

**Video 3. Three-dimensional model of a low-resolution spindle as used for the analysis of spindle geometry in a siMCRS1 cell**

Series of electron micrographs of a spindle in a siMCRS1 cell #5. The stacking of serial tomograms to generate a 3D model of the spindle volume is illustrated. The spindle on each section is indicated by a transparent gray area. Chromosomes are shown in blue. This video corresponds to the spindle shown in **Fig. 2D, right panel**.

**Video 4. Generation of a 3D model from joined serial electron tomograms displaying spindle siMCRS1 #1**

Series of tomograms and corresponding 3D model of spindle siMCRS1 #1. The stacking of serial tomograms to increase the tomographic volume is illustrated. Non-KMTs are shown as yellow lines, and KMTs are illustrated in red. This spindle reconstruction is not shown in any of the presented figures but has been added to show that the morphological alterations upon MCRS1-silencing are consistent between different 3D models. This video corresponds to spindle siMCRS1 #1 as given in **Table 1**.

**Video 5. Generation of a 3D model from joined serial electron tomograms displaying spindle siMCRS1 #5**

Series of tomograms and corresponding 3D model of spindle siMCRS1 #5. The stacking of serial tomograms to increase the tomographic volume is illustrated. KMTs are shown as red lines. This video corresponds to the spindle shown in **Fig. 3A**.

**Video 6. KMTs end morphology in spindle siScramble #2**

Three-dimensional model of KMTs with annotated end morphology. KMTs are indicated as black lines. Open KMT ends are labeled with green spheres, closed ends with purple spheres and undefined ends with white spheres. This video corresponds to the spindle shown in **Fig. 3A**

**Video 7: KMT end morphology in spindle siMCRS1 #5**

Three-dimensional model of KMTs with annotated end morphology. KMTs are indicated as black lines. Open KMT ends are labeled with green spheres, closed ends with purple spheres and undefined ends with white spheres. This video corresponds to the spindle shown in **Fig 3A.** and **Fig. 4F**.

## References

Akhmanova, A., and M.O. Steinmetz. 2019. Microtubule minus-end regulation at a glance. J Cell Sci. 132.

Barisic, M., G. Rajendraprasad, and Y. Steblyanko. 2021. The metaphase spindle at steady state - Mechanism and functions of microtubule poleward flux. Semin Cell Dev Biol. 117:99–117.

Brilot, A.F., A.S. Lyon, A. Zelter, S. Viswanath, A. Maxwell, M.J. MacCoss, E.G. Muller, A. Sali, T.N. Davis, and D.A. Agard. 2021. CM1-driven assembly and activation of yeast gamma-tubulin small complex underlies microtubule nucleation. Elife. 10.

Cheeseman, I.M., and A. Desai. 2008. Molecular architecture of the kinetochore-microtubule interface. Nat Rev Mol Cell Biol. 9:33–46.

Chen, X., L.A. Widmer, M.M. Stangier, M.O. Steinmetz, J. Stelling, and Y. Barral. 2019. Remote control of microtubule plus-end dynamics and function from the minus-end. Elife. 8.

Chretien, D., S.D. Fuller, and E. Karsenti. 1995. Structure of growing microtubule ends: two-dimensional sheets close into tubes at variable rates. J Cell Biol. 129:1311–1328.

Consolati, T., J. Locke, J. Roostalu, Z.A. Chen, J. Gannon, J. Asthana, W.M. Lim, F. Martino, M.A. Cvetkovic, J. Rappsilber, A. Costa, and T. Surrey. 2020. Microtubule Nucleation Properties of Single Human gammaTuRCs Explained by Their Cryo-EM Structure. Dev Cell. 53:603–617 e608.

DeLuca, J.G., W.E. Gall, C. Ciferri, D. Cimini, A. Musacchio, and E.D. Salmon. 2006. Kinetochore microtubule dynamics and attachment stability are regulated by Hec1. Cell. 127:969–982.

Detlev, S., M. Westerhoff, and H.-C. Hege. 2005. Amira: A Highly Interactive System for Visual Data Analysis “The visualization handbook.”.

Drutovic, D., X. Duan, R. Li, P. Kalab, and P. Solc. 2020. RanGTP and importin beta regulate meiosis I spindle assembly and function in mouse oocytes. EMBO J. 39:e101689.

Dudka, D., C. Castrogiovanni, N. Liaudet, H. Vassal, and P. Meraldi. 2019. Spindle-Length-Dependent HURP Localization Allows Centrosomes to Control Kinetochore-Fiber Plus-End Dynamics. Curr Biol. 29:3563–3578 e3566.

Elting, M.W., C.L. Hueschen, D.B. Udy, and S. Dumont. 2014. Force on spindle microtubule minus ends moves chromosomes. J Cell Biol. 206:245–256.

Ems-McClung, S.C., and C.E. Walczak. 2010. Kinesin-13s in mitosis: Key players in the spatial and temporal organization of spindle microtubules. Semin Cell Dev Biol. 21:276–282.

Foley, E.A., and T.M. Kapoor. 2013. Microtubule attachment and spindle assembly checkpoint signalling at the kinetochore. Nat Rev Mol Cell Biol. 14:25–37.

Ganem, N.J., and D.A. Compton. 2004. The KinI kinesin Kif2a is required for bipolar spindle assembly through a functional relationship with MCAK. J Cell Biol. 166:473–478.

Ganem, N.J., and D.A. Compton. 2006. Functional roles of poleward microtubule flux during mitosis. Cell Cycle. 5:481–485.

Ganem, N.J., K. Upton, and D.A. Compton. 2005. Efficient mitosis in human cells lacking poleward microtubule flux. Curr Biol. 15:1827–1832.

Gudimchuk, N.B., and J.R. McIntosh. 2021. Regulation of microtubule dynamics, mechanics and function through the growing tip. Nat Rev Mol Cell Biol. 22:777–795.

Gudimchuk, N.B., E.V. Ulyanov, E. O’Toole, C.L. Page, D.S. Vinogradov, G. Morgan, G. Li, J.K. Moore, E. Szczesna, A. Roll-Mecak, F.I. Ataullakhanov, and J. Richard McIntosh. 2020. Mechanisms of microtubule dynamics and force generation examined with computational modeling and electron cryotomography. Nat Commun. 11:3765.

Hoog, J.L., S.M. Huisman, Z. Sebo-Lemke, L. Sandblad, J.R. McIntosh, C. Antony, and D. Brunner. 2011. Electron tomography reveals a flared morphology on growing microtubule ends. J Cell Sci. 124:693–698.

Jiang, K., L. Rezabkova, S. Hua, Q. Liu, G. Capitani, A.F.M. Altelaar, A.J.R. Heck, R.A. Kammerer, M.O. Steinmetz, and A. Akhmanova. 2017. Microtubule minus-end regulation at spindle poles by an ASPM-katanin complex. Nat Cell Biol. 19:480–492.

Kamasaki, T., E. O’Toole, S. Kita, M. Osumi, J. Usukura, J.R. McIntosh, and G. Goshima. 2013. Augmin-dependent microtubule nucleation at microtubule walls in the spindle. J Cell Biol. 202:25–33.

Keating, T.J., and G.G. Borisy. 2000. Immunostructural evidence for the template mechanism of microtubule nucleation. Nat Cell Biol. 2:352–357.

Kiewisz, R., G. Fabig, W. Conway, D. Baum, D. Needleman, and T. Muller-Reichert. 2022. Three-dimensional structure of kinetochore-fibers in human mitotic spindles. Elife. 11.

Kiewisz, R., T. Muller-Reichert, and G. Fabig. 2021. High-throughput screening of mitotic mammalian cells for electron microscopy using classic histological dyes. Methods Cell Biol. 162:151–170.

Kirschner, M.W., and T. Mitchison. 1986. Microtubule dynamics. Nature. 324:621.

Kollman, J.M., C.H. Greenberg, S. Li, M. Moritz, A. Zelter, K.K. Fong, J.J. Fernandez, A. Sali, J. Kilmartin, T.N. Davis, and D.A. Agard. 2015. Ring closure activates yeast gammaTuRC for species-specific microtubule nucleation. Nat Struct Mol Biol. 22:132–137.

Lampson, M.A., and E.L. Grishchuk. 2017. Mechanisms to Avoid and Correct Erroneous Kinetochore-Microtubule Attachments. Biology (Basel). 6.

Lindow, N., F.N. Brunig, V.J. Dercksen, G. Fabig, R. Kiewisz, S. Redemann, T. Muller-Reichert, S. Prohaska, and D. Baum. 2021. Semi-automatic stitching of filamentous structures in image stacks from serial-section electron tomography. J Microsc. 284:25–44.

Lüders, J. 2016. The Microtubule Cytoskeleton: Organisation, Function and Role in Disease. The Microtubule Cytoskeleton: Organisation, Function and Role in Disease. Springer.

Maiato, H., J. DeLuca, E.D. Salmon, and W.C. Earnshaw. 2004. The dynamic kinetochore-microtubule interface. J Cell Sci. 117:5461–5477.

Mandelkow, E.M., E. Mandelkow, and R.A. Milligan. 1991. Microtubule dynamics and microtubule caps: a time-resolved cryo-electron microscopy study. J Cell Biol. 114:977–991.

Maresca, T.J., and E.D. Salmon. 2009. Intrakinetochore stretch is associated with changes in kinetochore phosphorylation and spindle assembly checkpoint activity. J Cell Biol. 184:373–381.

Mastronarde, D.N. 1997. Dual-axis tomography: an approach with alignment methods that preserve resolution. J Struct Biol. 120:343–352.

Mastronarde, D.N. 2003. SerialEM: A Program for Automated Tilt Series Acquisition on Tecnai Microscopes Using Prediction of Specimen Position. Micros Microanal. 9:1182–| 1183.

Mastronarde, D.N. 2005. Automated electron microscope tomography using robust prediction of specimen movements. J Struct Biol. 152:36–51.

Mastronarde, D.N., and S.R. Held. 2017. Automated tilt series alignment and tomographic reconstruction in IMOD. J Struct Biol. 197:102–113.

McEwen, B.F., and M. Marko. 1999. Three-dimensional transmission electron microscopy and its application to mitosis research. Methods Cell Biol. 61:81–111.

McIntosh, J.R., E.L. Grishchuk, M.K. Morphew, A.K. Efremov, K. Zhudenkov, V.A. Volkov, I.M. Cheeseman, A. Desai, D.N. Mastronarde, and F.I. Ataullakhanov. 2008. Fibrils connect microtubule tips with kinetochores: a mechanism to couple tubulin dynamics to chromosome motion. Cell. 135:322–333.

McIntosh, J.R., E. O’Toole, G. Morgan, J. Austin, E. Ulyanov, F. Ataullakhanov, and N. Gudimchuk. 2018. Microtubules grow by the addition of bent guanosine triphosphate tubulin to the tips of curved protofilaments. J Cell Biol. 217:2691–2708.

McIntosh, J.R., E. O’Toole, K. Zhudenkov, M. Morphew, C. Schwartz, F.I. Ataullakhanov, and E.L. Grishchuk. 2013. Conserved and divergent features of kinetochores and spindle microtubule ends from five species. J Cell Biol. 200:459–474.

Meunier, S., M. Shvedunova, N. Van Nguyen, L. Avila, I. Vernos, and A. Akhtar. 2015. An epigenetic regulator emerges as microtubule minus-end binding and stabilizing factor in mitosis. Nat Commun. 6:7889.

Meunier, S., and I. Vernos. 2011. K-fibre minus ends are stabilized by a RanGTP-dependent mechanism essential for functional spindle assembly. Nat Cell Biol. 13:1406–1414.

Meunier, S., and I. Vernos. 2012. Microtubule assembly during mitosis - from distinct origins to distinct functions? J Cell Sci. 125:2805–2814.

Meunier, S., and I. Vernos. 2016. Acentrosomal Microtubule Assembly in Mitosis: The Where, When, and How. Trends Cell Biol. 26:80–87.

Mimori-Kiyosue, Y., and S. Tsukita. 2003. “Search-and-capture” of microtubules through plus-end-binding proteins (+TIPs). J Biochem. 134:321–326.

Mitchison, T., and M. Kirschner. 1984. Dynamic instability of microtubule growth. Nature. 312:237–242.

Mitchison, T.J. 1989. Polewards microtubule flux in the mitotic spindle: evidence from photoactivation of fluorescence. J Cell Biol. 109:637–652.

Monda, J.K., and I.M. Cheeseman. 2018. The kinetochore-microtubule interface at a glance. J Cell Sci. 131.

Moritz, M., M.B. Braunfeld, V. Guenebaut, J. Heuser, and D.A. Agard. 2000. Structure of the gamma-tubulin ring complex: a template for microtubule nucleation. Nat Cell Biol. 2:365–370.

Muller-Reichert, T., D. Chretien, F. Severin, and A.A. Hyman. 1998. Structural changes at microtubule ends accompanying GTP hydrolysis: information from a slowly hydrolyzable analogue of GTP, guanylyl (alpha,beta)methylenediphosphonate. Proc Natl Acad Sci U S A. 95:3661–3666.

Muller-Reichert, T., H. Hohenberg, E.T. O’Toole, and K. McDonald. 2003. Cryoimmobilization and three-dimensional visualization of C. elegans ultrastructure. J Microsc. 212:71–80.

O’Toole, E., M. Morphew, and J.R. McIntosh. 2020. Electron tomography reveals aspects of spindle structure important for mechanical stability at metaphase. Mol Biol Cell. 31:184–195.

O’Toole, E.T., K.L. McDonald, J. Mantler, J.R. McIntosh, A.A. Hyman, and T. Muller-Reichert. 2003. Morphologically distinct microtubule ends in the mitotic centrosome of Caenorhabditis elegans. J Cell Biol. 163:451–456.

Redemann, S., J. Baumgart, N. Lindow, M. Shelley, E. Nazockdast, A. Kratz, S. Prohaska, J. Brugues, S. Furthauer, and T. Muller-Reichert. 2017. C. elegans chromosomes connect to centrosomes by anchoring into the spindle network. Nat Commun. 8:15288.

Redemann, S., B. Weber, M. Moller, J.M. Verbavatz, A.A. Hyman, D. Baum, S. Prohaska, and T. Muller-Reichert. 2014. The segmentation of microtubules in electron tomograms using Amira. Methods Mol Biol. 1136:261–278.

Rice, L.M. 2018. A new look for the growing microtubule end? J Cell Biol. 217:2609–2611.

Simon, J.R., and E.D. Salmon. 1990. The structure of microtubule ends during the elongation and shortening phases of dynamic instability examined by negative-stain electron microscopy. J Cell Sci. 96 (Pt 4):571–582.

Steblyanko, Y., G. Rajendraprasad, M. Osswald, S. Eibes, A. Jacome, S. Geley, A.J. Pereira, H. Maiato, and M. Barisic. 2020. Microtubule poleward flux in human cells is driven by the coordinated action of four kinesins. EMBO J. 39:e105432.

Teixido-Travesa, N., J. Roig, and J. Luders. 2012. The where, when and how of microtubule nucleation - one ring to rule them all. J Cell Sci. 125:4445–4456.

VandenBeldt, K.J., R.M. Barnard, P.J. Hergert, X. Meng, H. Maiato, and B.F. McEwen. 2006. Kinetochores use a novel mechanism for coordinating the dynamics of individual microtubules. Curr Biol. 16:1217–1223.

Waters, J.C., T.J. Mitchison, C.L. Rieder, and E.D. Salmon. 1996. The kinetochore microtubule minus-end disassembly associated with poleward flux produces a force that can do work. Mol Biol Cell. 7:1547–1558.

Weber, B., G. Greenan, S. Prohaska, D. Baum, H.C. Hege, T. Muller-Reichert, A.A. Hyman, and J.M. Verbavatz. 2012. Automated tracing of microtubules in electron tomograms of plastic embedded samples of Caenorhabditis elegans embryos. J Struct Biol. 178:129–138.

Weber, B., E.M. Tranfield, J.L. Hoog, D. Baum, C. Antony, T. Hyman, J.M. Verbavatz, and S. Prohaska. 2014. Automated stitching of microtubule centerlines across serial electron tomograms. PLoS One. 9:e113222.

Wieczorek, M., L. Urnavicius, S.C. Ti, K.R. Molloy, B.T. Chait, and T.M. Kapoor. 2020. Asymmetric Molecular Architecture of the Human gamma-Tubulin Ring Complex. Cell. 180:165–175 e116.

Wiese, C., and Y. Zheng. 2000. A new function for the gamma-tubulin ring complex as a microtubule minus-end cap. Nat Cell Biol. 2:358–364.

Zheng, Y., M.L. Wong, B. Alberts, and T. Mitchison. 1995. Nucleation of microtubule assembly by a gamma-tubulin-containing ring complex. Nature. 378:578–583.

Zimmermann, F., M. Serna, A. Ezquerra, R. Fernandez-Leiro, O. Llorca, and J. Luders. 2020. Assembly of the asymmetric human gamma-tubulin ring complex by RUVBL1-RUVBL2 AAA ATPase. Sci Adv. 6.

