## Supplementary figures and images for "MCRS1 modulates the heterogeneity of microtubule minus-end morphologies in mitotic spindles"

### Figure S1

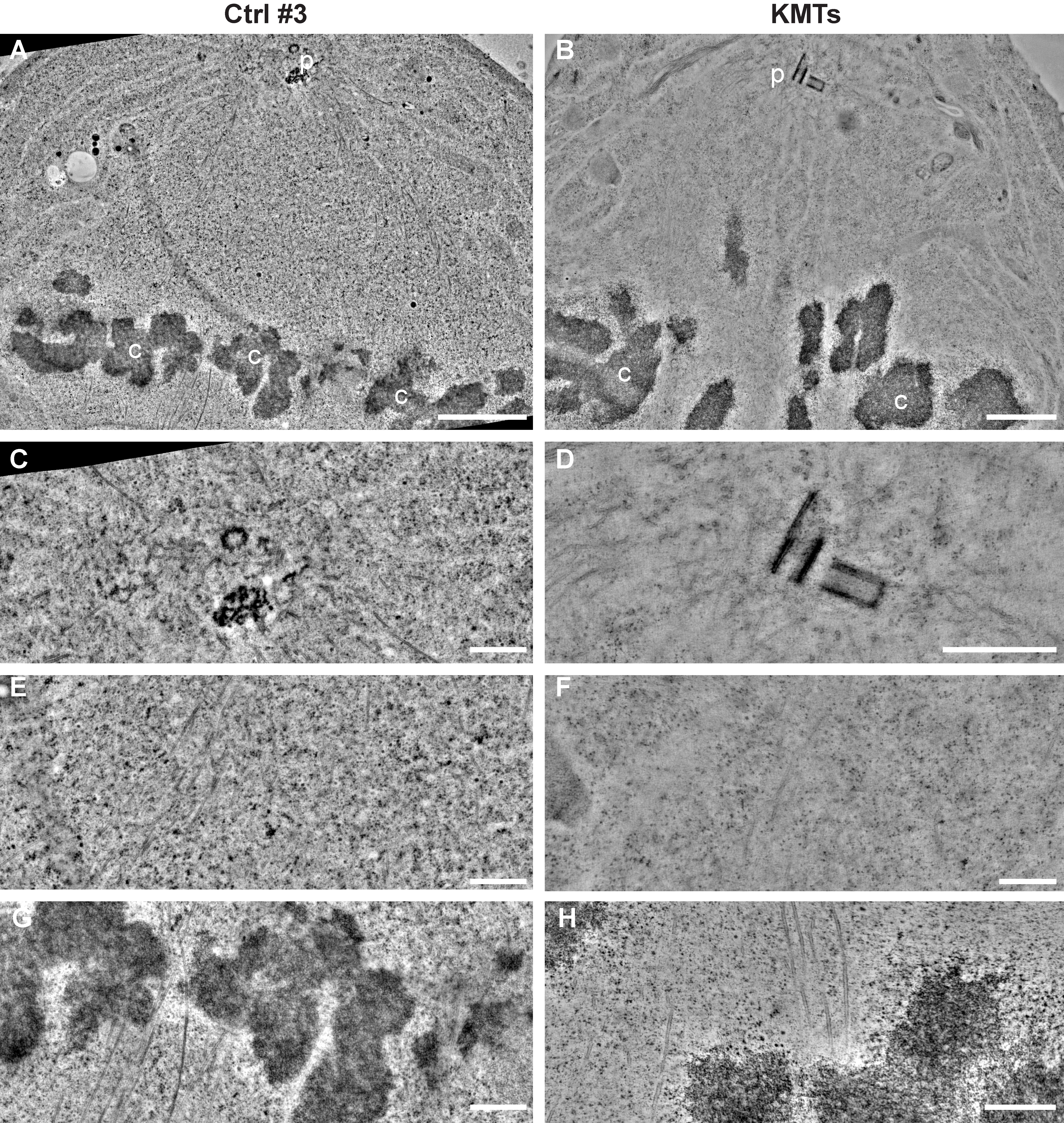

### Figure S2

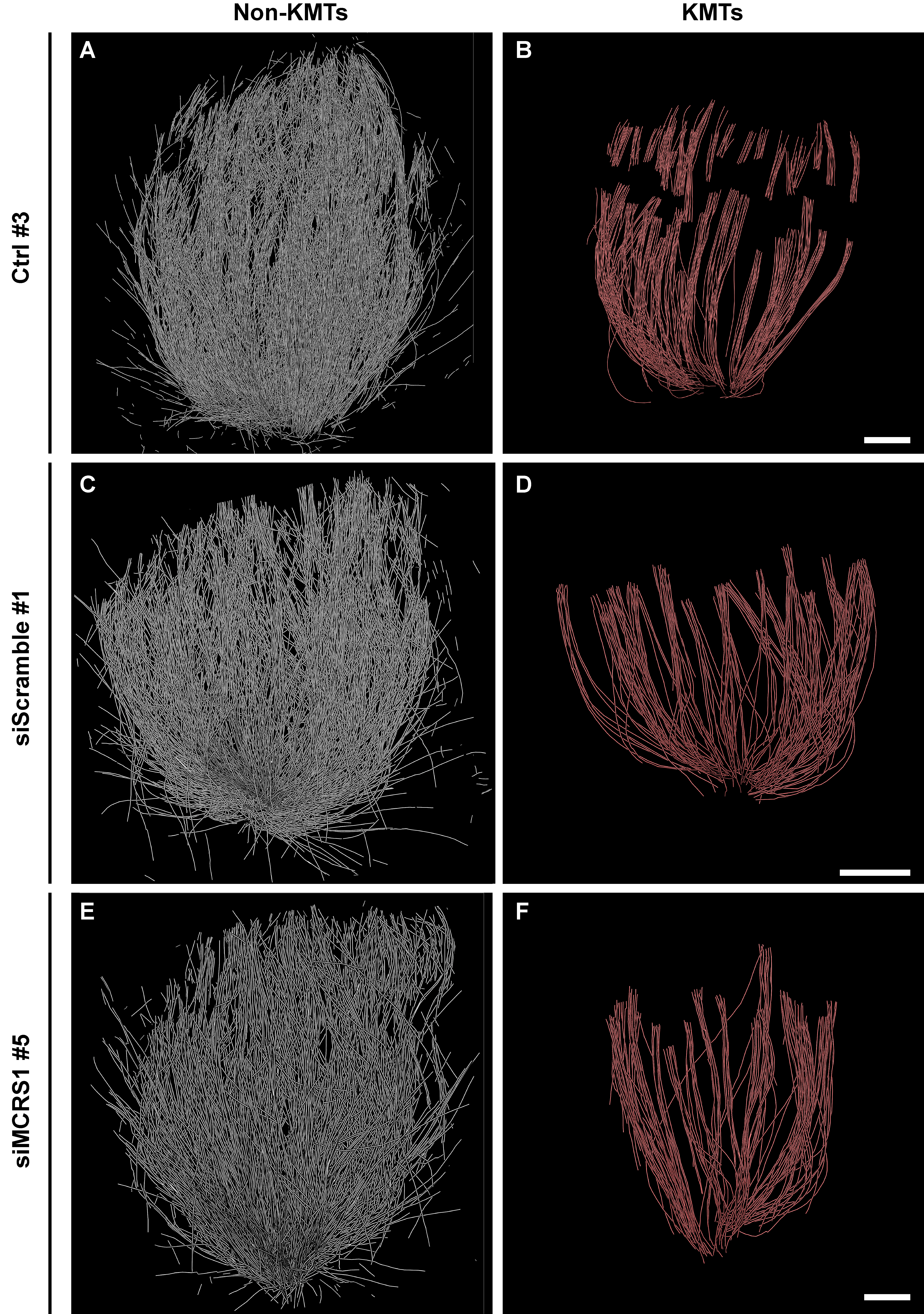

### Figure S3

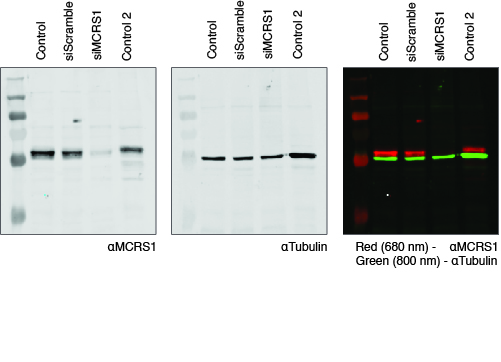

### Figure S4

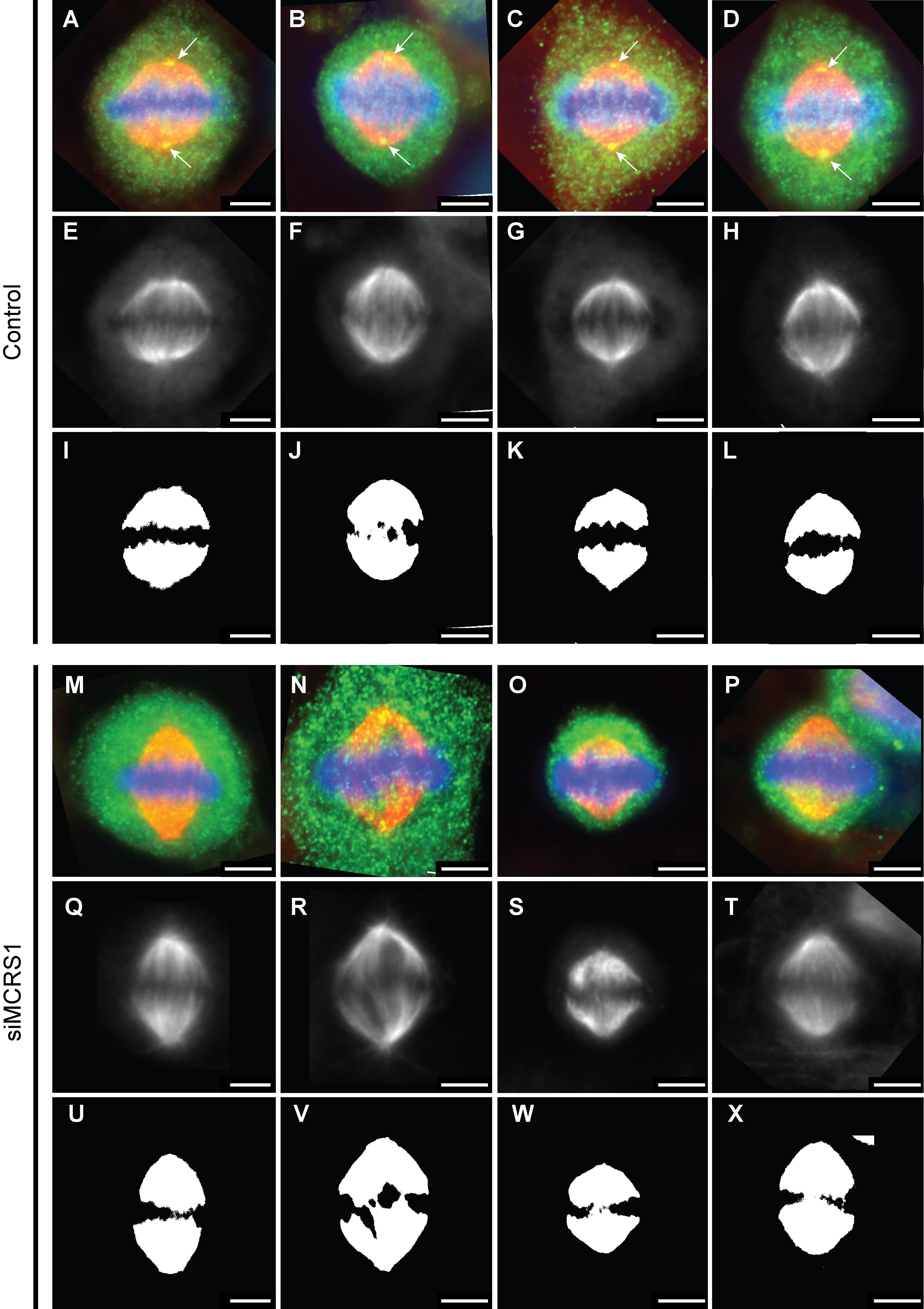

### Figure S5

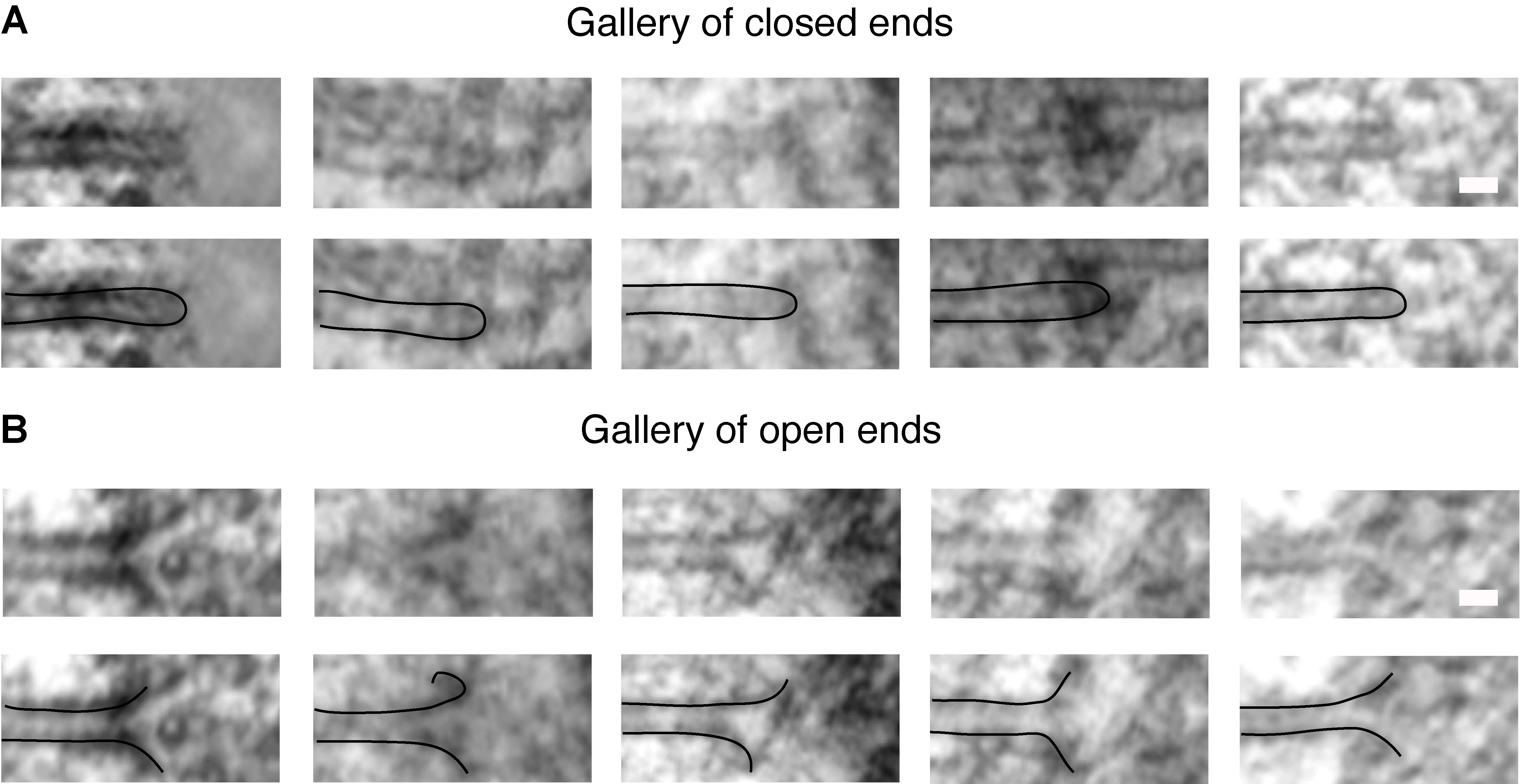

### Figure S6

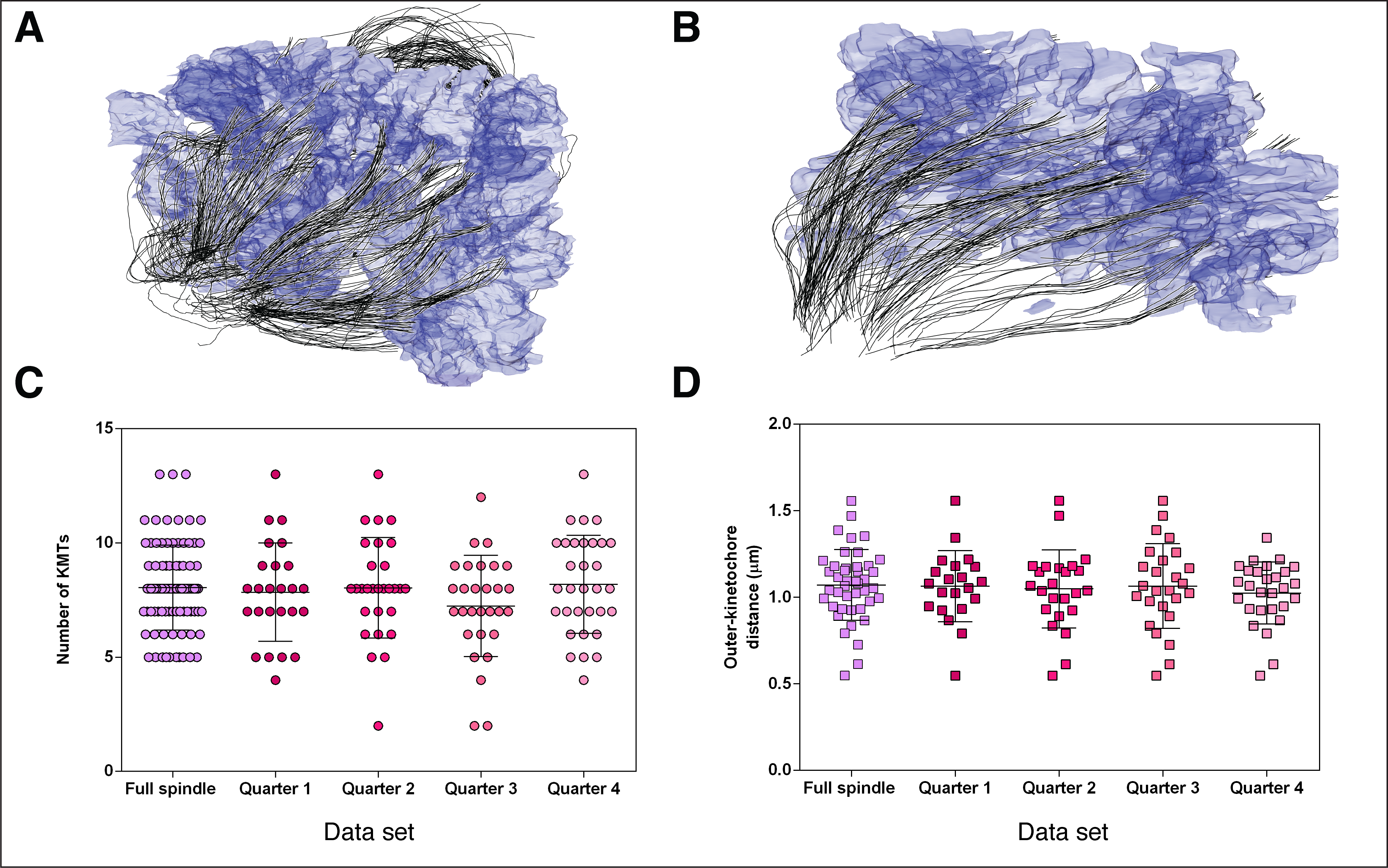

### Figure S7

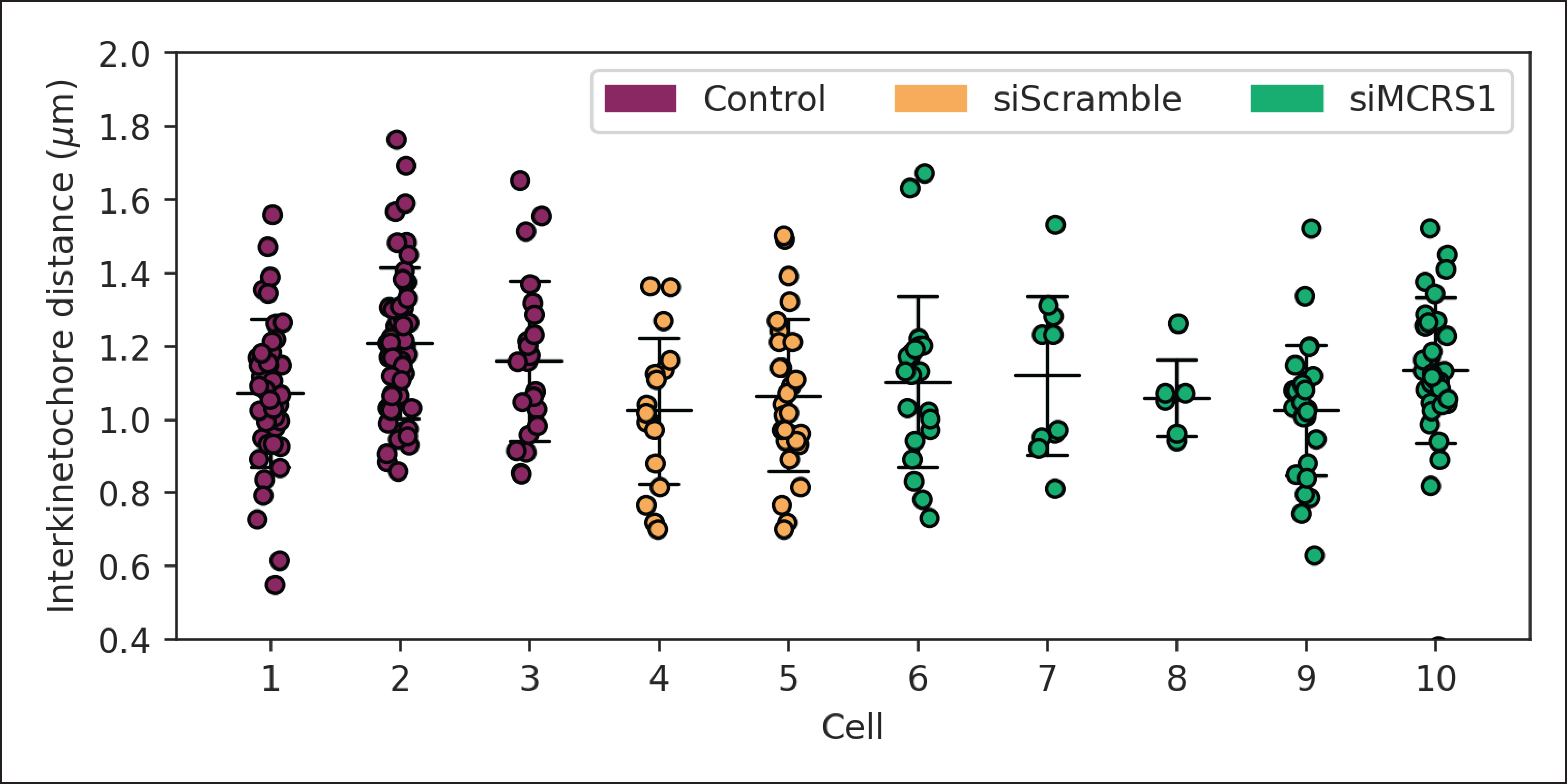

### Figure S8

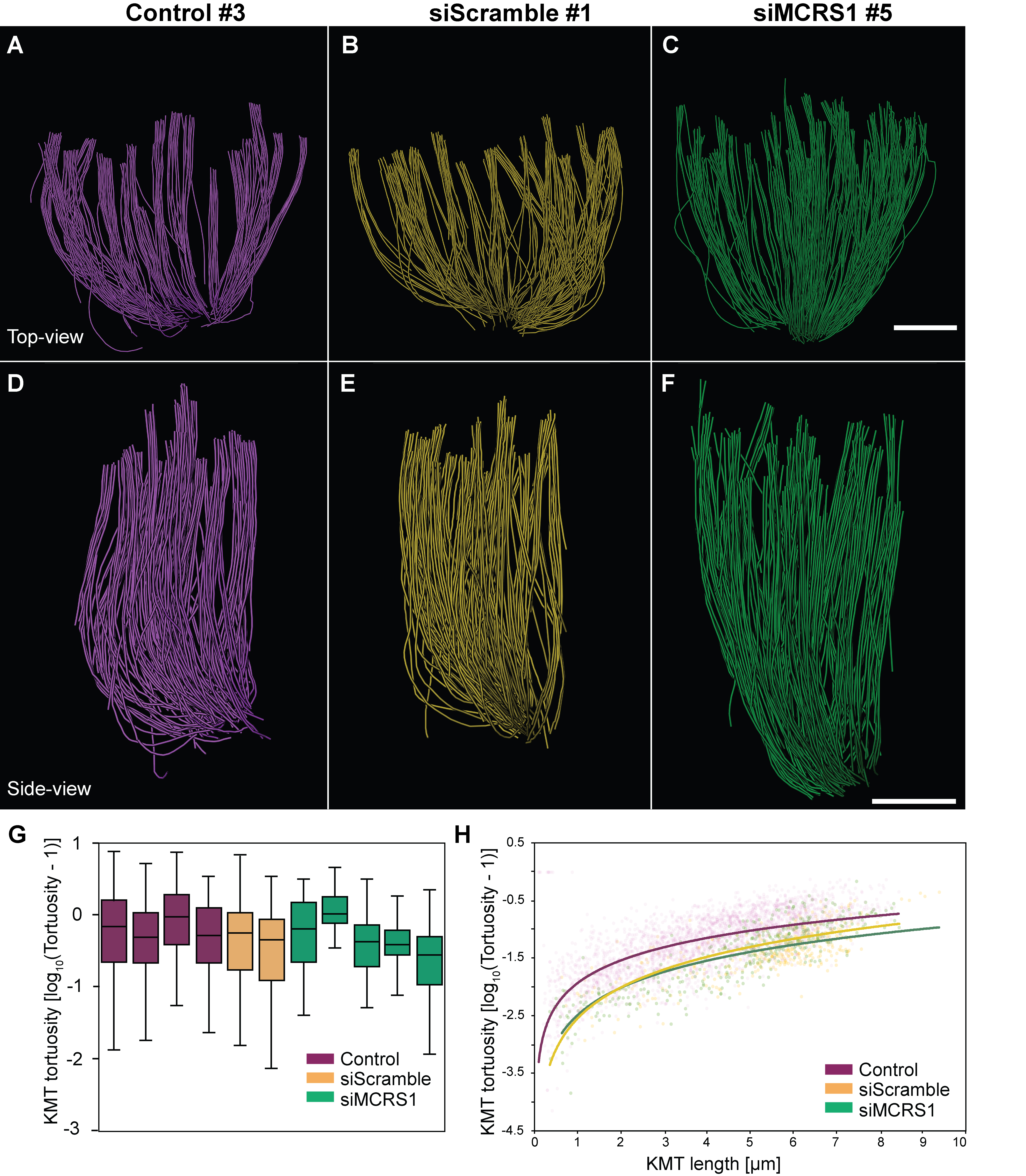
